# CDK19 is a Regulator of Triple-Negative Breast Cancer Growth

**DOI:** 10.1101/317776

**Authors:** Robert W. Hsieh, Angera H. Kuo, Ferenc A. Scheeren, Mark A. Zarnegar, Shaheen S. Sikandar, Jane Antony, Luuk S. Heitink, Divya Periyakoil, Tomer Kalisky, Sopheak Sim, Dalong Qian, Sanjay V. Malhotra, George Somlo, Frederick M. Dirbas, Ajit Jadhav, Aaron M. Newman, Michael F. Clarke

**Affiliations:** Institute for Stem Cell Biology and Regenerative Medicine, School of Medicine, Stanford University, Stanford, CA 94305, USA; Division of Oncology and Stanford Cancer Institute, School of Medicine, Stanford University, Stanford, CA 94305, USA; Department of Medical Oncology, Department of Surgery and Immunohematology and Blood Transfusion, Leiden University Medical Center (LUMC), Leiden, the Netherlands.; Faculty of Engineering, Bar-Ilan University, Ramat Gan, 52900, Israel; Division of Radiation Oncology, School of Medicine, Stanford University, Stanford, CA 94305, USA.; Department of Medical Oncology, Beckman Research Institute and City of Hope Comprehensive Cancer Center, Duarte, CA 91010, USA.; Division of Surgical Oncology, Department of Surgery, School of Medicine, Stanford University, Stanford, CA 94305, USA.; National Center for Advancing Translational Sciences (NCATS), National Institute of Health, Bethesda, MD 20892, USA.; Chan Zuckerberg Biohub Investigator

## Abstract

Triple-negative breast cancer (TNBC) is a poor prognosis disease with no clinically approved targeted therapies. Here, using *in vitro* and *in vivo* RNA interference (RNAi) screens in TNBC patient-derived xenografts (PDX), we identify cyclin dependent kinase 19 (CDK19) as a potential therapeutic target. Using *in vitro* and *in vivo* TNBC PDX models, we validated the inhibitory effect of *CDK19* knockdown on tumor initiation, proliferation and metastases. Despite this, *CDK19* knockdown did not affect the growth of non-transformed mammary epithelial cells. Using CD10 and EpCAM as novel tumor initiating cell (TIC) markers, we found the EpCAM^med/high^/CD10^−/low^ TIC sub-population to be enriched in CDK19 and a putative cellular target of *CDK19* inhibition. Comparative gene expression analysis of *CDK19* and *CDK8* knockdowns revealed that CDK19 regulates a number of cancer-relevant pathways, uniquely through its own action and others in common with CDK8. Furthermore, although it is known that CDK19 can act at enhancers, our CHIP-Seq studies showed that CDK19 can also epigenetically modulate specific H3K27Ac enhancer signals which correlate with gene expression changes. Finally, to assess the potential therapeutic utility of CDK19, we showed that both *CDK19* knockdown and chemical inhibition of CDK19 kinase activity impaired the growth of pre-established PDX tumors *in vivo*. Current strategies inhibiting transcriptional co-factors and targeting TICs have been limited by toxicity to normal cells. Because of CDK19’s limited tissue distribution and the viability of *CDK19* knockout mice, CDK19 represents a promising therapeutic target for TNBC.

## Introduction

Cyclin Dependent Kinase (CDK) 19 and its paralog, CDK8, are members of the transcriptional CDKs which also include CDK7, CDK9, CDK12 and CDK13. Unlike other CDKs, these transcriptional CDKs are less involved in cell cycle regulatory processes and are more involved in transcription^1^. CDK19 and its paralog, CDK8, both interact with Cyclin C as well as Mediator 12 and Mediator 13 (or their paralogs Mediator 12L and Mediator 13L) to form a CDK module (CKM)^2,3^. In turn, the CKM interacts with core Mediator to regulate RNA polymerase II (Pol II) association and transcriptional activity.

CDK8, in particular, has been a focus of intense study as it has been implicated in in a number of important pathways such as WNT/beta-catenin signaling, KRAS and Notch in a number of cancers including colon cancer^4^, acute myeloid leukemia^5^ and melanoma^6^. In contrast, much less is known about CDK19. Initially, CDK19 was thought to function similarly or redundantly to CDK8 given that they share 84% amino acid sequence similarity (Supplementary Fig. 1A) and 97% identity in the kinase domain^7^. However, functional differences have since been uncovered. Recent, *in vitro* studies showed that CDK19 and CDK8 participate mutually exclusively of each other in binding to other CKM components^8^, while gene knockdown studies in cell lines of cervical cancer^9^ and colon cancer^8^ showed that both CDK19 and CDK8 can regulate different genes.

Overall, CDK19 also appears to be more specialized in its role than CDK8. Evolutionarily, CDK8 is expressed in all eukaryotes, whereas CDK19 is expressed only in vertebrates. Also, compared to the ubiquitous expression of CDK8, CDK19 has a much more limited tissue distribution^9^, thus offering the theoretical benefit of less systemic toxicity from CDK19 inhibition compared to CDK8 inhibition. Further support of this potential benefit is the finding that in the normal human colon^10^, a common site of dose limiting drug toxicity, there is much lower expression of *CDK19* than *CDK8* (Supplementary Fig. 1B). It is also worth noting that *CDK8* knockouts are lethal^11^ whereas *CDK19* knockout mice are viable^12^. Collectively, the data provides evidence to suggest that CDK19 may potentially be a superior therapeutic target compared to CDK8.

We show here that CDK19, identified from two pooled RNAi dropout viability screens and subsequently validated comprehensively *in vitro* and *in vivo*, is important to the growth of TNBC. As patients with advanced TNBC currently have poor overall prognoses^13^ and lack clinically approved targeted therapies, there is an urgent medical need to investigate potential therapeutic targets, such as CDK19, for TNBC treatment.

## Results

### CDK19 was identified from two RNAi dropout viability screens as important to the growth of TNBC

To identify genes essential for the growth of TNBC, we performed two pooled RNAi dropout viability screens using a 27,500 shRNA library targeting 5000 genes^14^ in PDX-T1^15^, a TNBC PDX (Supplementary Table 1). The screens were performed in two different formats, *in vitro* as organoid cultures and *in vivo* as PDXs in *nod scid gamma* (NSG) mice (Supplementary Fig. 2A). The goal was to identify genes whose knockdown by shRNA inhibited the growth of PDX tumor cells across different experimental conditions. We restricted our final candidate list to genes with the lowest 5% of shRNA ratios in each screen that were targeted by more than two shRNAs and were also identified both *in vitro* and *in vivo* (Supplementary Fig. 2B). This resulted in the identification of 46 candidate genes (Supplementary Fig. 2C).

CDK19 was selected to pursue further because of evidence suggesting that it has biological and clinical relevance to TNBC. From The Cancer Genome Atlas (TCGA)^16^ data we found that *CDK19* copy number amplifications and mRNA upregulation were more prevalent in TNBC patient samples (16%) compared to samples from other breast cancer subtypes (Supplementary Fig. 2D). Of clinical relevance, high *CDK19* expression has been reported to correlate with poor relapse free survival in *all* breast cancer patients^17,18^. However, more compelling is the finding that there is a statistically significant association of *CDK19* expression with relapse free survival in the TNBC subtype (this is also true of the ER+ subtype, but not the HER2+ subtype) (Supplementary Fig. 2E).

### Constitutive *CDK19* knockdown suppresses the *in vitro* growth of TNBC in both cell lines and organoids but not in non-transformed human mammary epithelial cells

To validate the growth inhibitory effect of *CDK19* knockdown in TNBC, we initially utilized two commonly used TNBC cell lines: MDA-MB231 and MDA-MB468. Using two different constitutively expressed shRNAs (shCDK19-1 and shCDK19-2) targeting *CDK19*, we confirmed knockdown of *CDK19* (Supplementary Fig. 3A and 3B), and showed that it decreased cell proliferation in both TNBC cell lines (Fig. 1A and 1B). In a more complex *in vitro* model with patient-derived xenograft (PDX) tumors growing as organoids, *CDK19* knockdown (Supplementary Fig. 3C) also inhibited the formation of TNBC PDX-T1 organoid colonies (Fig. 1C). To determine the effects of *CDK19* knockdown in non-transformed mammary cells, we infected human mammary epithelial cells (HMEC) with shRNA targeting *CDK19*. In HMECs, *CDK19* knockdown did not affect the viability of the cells (Fig. 1D). Collectively, our studies showed *in vitro* that *CDK19* knockdown inhibited the proliferation of TNBC cell lines and the formation of TNBC PDX organoid colonies but did not adversely affect the growth of non-transformed mammary epithelial cells.

**Figure 1.**
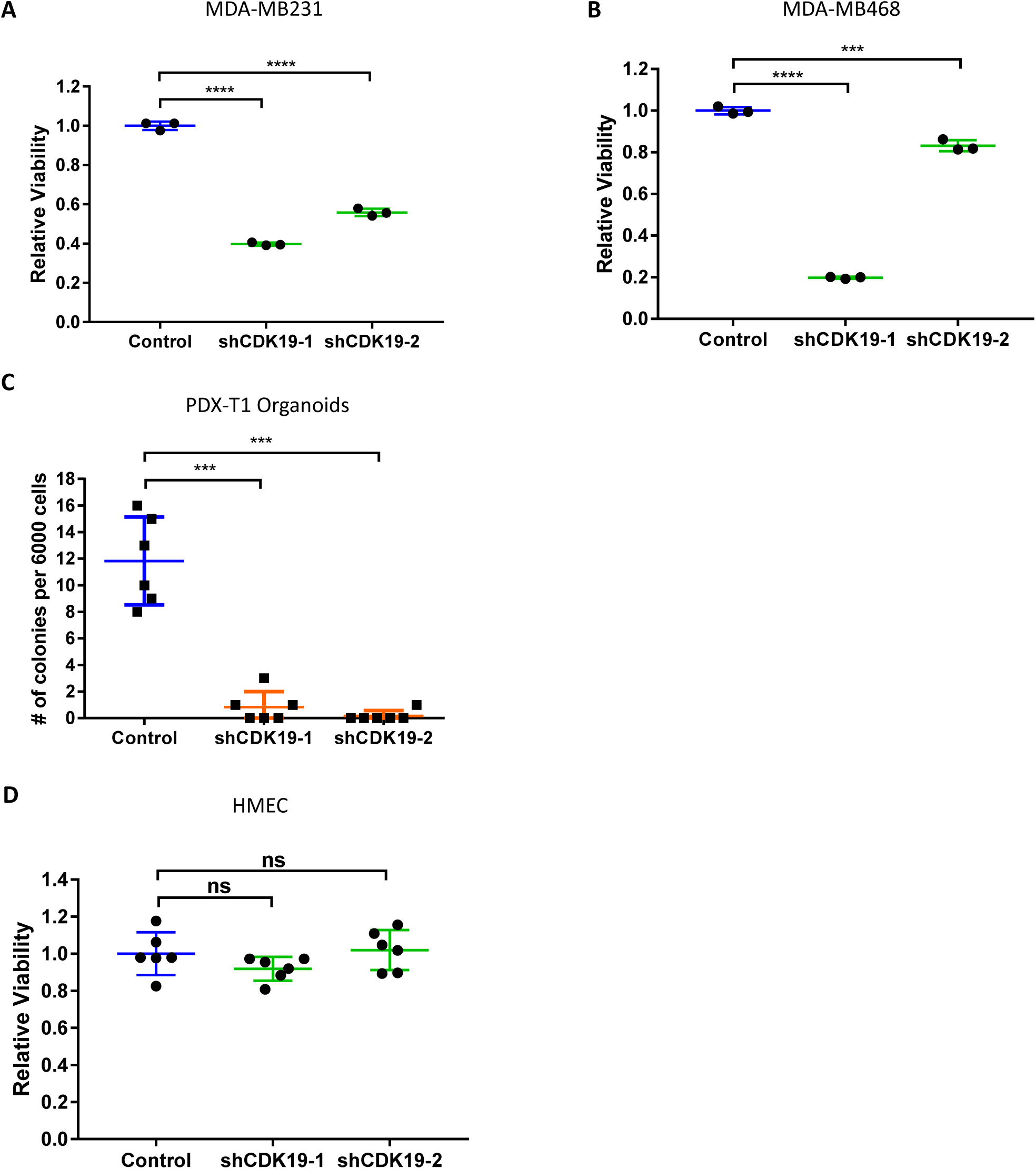
*CDK19* knockdown inhibits the growth of TNBC cells *in vitro* but does not affect the growth of normal mammary epithelial cells. (A-B) Viability of (A) MDA-MB231 cells and (B) MDA-MB468 cells assessed 4 days after transduction with control shRNA or *CDK19* targeting shRNA (shCDK19-1, shCDK19-2). All values were normalized to control shRNA sample (n = 3 experiment performed three times),.
(C) Number of organoid colonies formed two weeks after transduction of PDX cells with either control shRNA or *CDK19* targeting shRNA (shCDK19-1, shCDK19-2) (n = 6, experiment performed two times).
(D) Viability of HMEC cells assessed four days after transduction with control shRNA or *CDK19* targeting shRNA (shCDK19-1, shCDK19-2). All values normalized to the control shRNA sample (n = 6, experiment performed three times).

All data are represented as mean ± s.d., *P*-values determined by unpaired t-test. ns is *P* > 0.05, \*\*\**P* < 0.001, \*\*\*\**P* < 0.0001.

### Constitutive *CDK19* knockdown suppresses the *in vivo* growth of PDX tumors and inhibits tumor metastases

We extended our studies to more physiologically relevant *in vivo* models by knocking down *CDK19* in four different TNBC PDXs grown in *NOD scid gamma* (NSG) mice. Three of these PDXs: PDX-T1, PDX-T2^15^ and PDX-T3 were derived from chemotherapy naive patients (Supplementary Table 1). A fourth, PDX-T4, was an aggressive PDX obtained from the brain metastasis of a patient with a chemotherapy-resistant inflammatory breast cancer (Supplementary Table 1). Inflammatory breast cancers are known to be aggressive, difficult to treat and associated with extremely poor prognoses^19^. In PDX-T1, knockdown of *CDK19* impaired tumor growth (Fig. 2A). Due to the difficulty when measuring tumor volumes to differentiate between the growth of shRNA transduced cells and untransduced cells within the tumor, all PDX tumor cells were labeled with green fluorescent protein (GFP) and cells subsequently infected with either *CDK19* shRNA or control shRNA were additionally labeled with red fluorescent protein (RFP). By monitoring the change in RFP positive cells in the GFP-labelled tumor cells, we can monitor the effect of *CDK19* knockdown specifically in the transduced PDX tumor cells. Our studies showed that *CDK19* knockdown led to a significant reduction in the percentage of RFP positive cells in tumors from all three TNBC PDXs (Fig. 2B and 2C, 2D and 2E). Notably, this effect was even seen in PDX-T4 (Fig. 2F), which we anticipated to be least likely to respond due to its chemo-resistance. These results showed that CDK19 is also critical *in vivo* for tumor growth.

**Figure 2.**
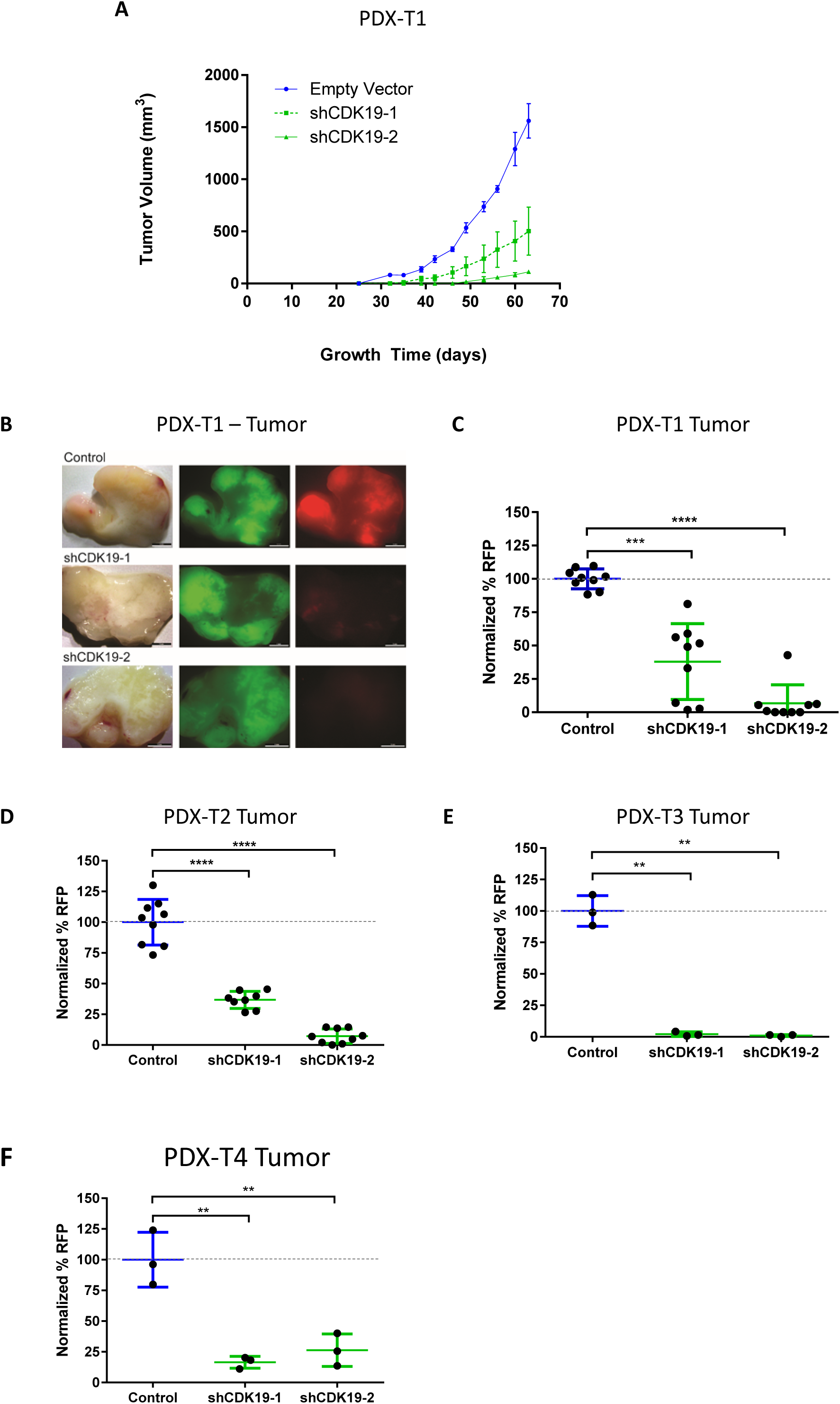
*CDK19* knockdown inhibits the growth of TNBC PDXs in NSG mice. (A) Representative plots showing volumes of PDX-T1 tumors transduced with control shRNA (blue line), shCDK19-1 (green dotted line) or shCDK19-2 (green solid line) as a function of time in NSG mice (mean ± s.d., 3 mice in each group, experiment performed three times).
(B) Representative images of PDX-T1 tumors transduced with control shRNA (top row), shCDK19-1 (middle row) or shCDK19-2 (bottom row). Brightfield images (left column) show gross tumor morphology, FITC images (middle column) identify tumor cells labeled with GFP (green) and Texas-Red images (right column) identify shRNA-transduced cells labeled with RFP (red). Scale bars = 5 mm.
(C-F) Flow cytometry analysis of RFP positivity in (C) PDX-T1, (D) PDX-T2, (E) PDX-T3 or (F) PDX-T4 tumor cells that were transduced with control shRNA or *CDK19* targeting shRNA (shCDK19-1, shCDK19-2) and grown in NSG mice. All final RFP percentages were normalized to the initial RFP infection percentages (to account for infection differences) and then normalized to the mean RFP percentage of the control shRNA sample. Each data point represents one mouse (mean ± s.d., PDX-T1 and PDX-T2 had 9 mice for each condition tested, PDX-T3 and PDX-T4 had 3 mice for each condition tested).

For all, *P*-values determined by unpaired t-test. \*\**P* < 0.01, \*\*\**P* < 0.001; \*\*\*\**P* < 0.0001.

Two of our PDX tumors, PDX-T1 and PDX-T4, routinely metastasize to the lungs of the host NSG mice. Remarkably, *CDK19* knockdown with either shRNA eliminated the metastases of transduced (RFP positive) tumor cells in both PDX-T1 (Fig. 3A and 3C) and PDX-T4 tumors (Fig. 3B and 3D). The lack of metastases in the *CDK19* knockdown groups were not attributable to differences in primary tumor volume (Supplementary Fig. 3E) or differences in the time since appearance of the primary tumor (Supplementary Fig. 3F). Thus, *in vivo*, *CDK19* knockdown also inhibited tumor metastasis.

**Figure 3.**
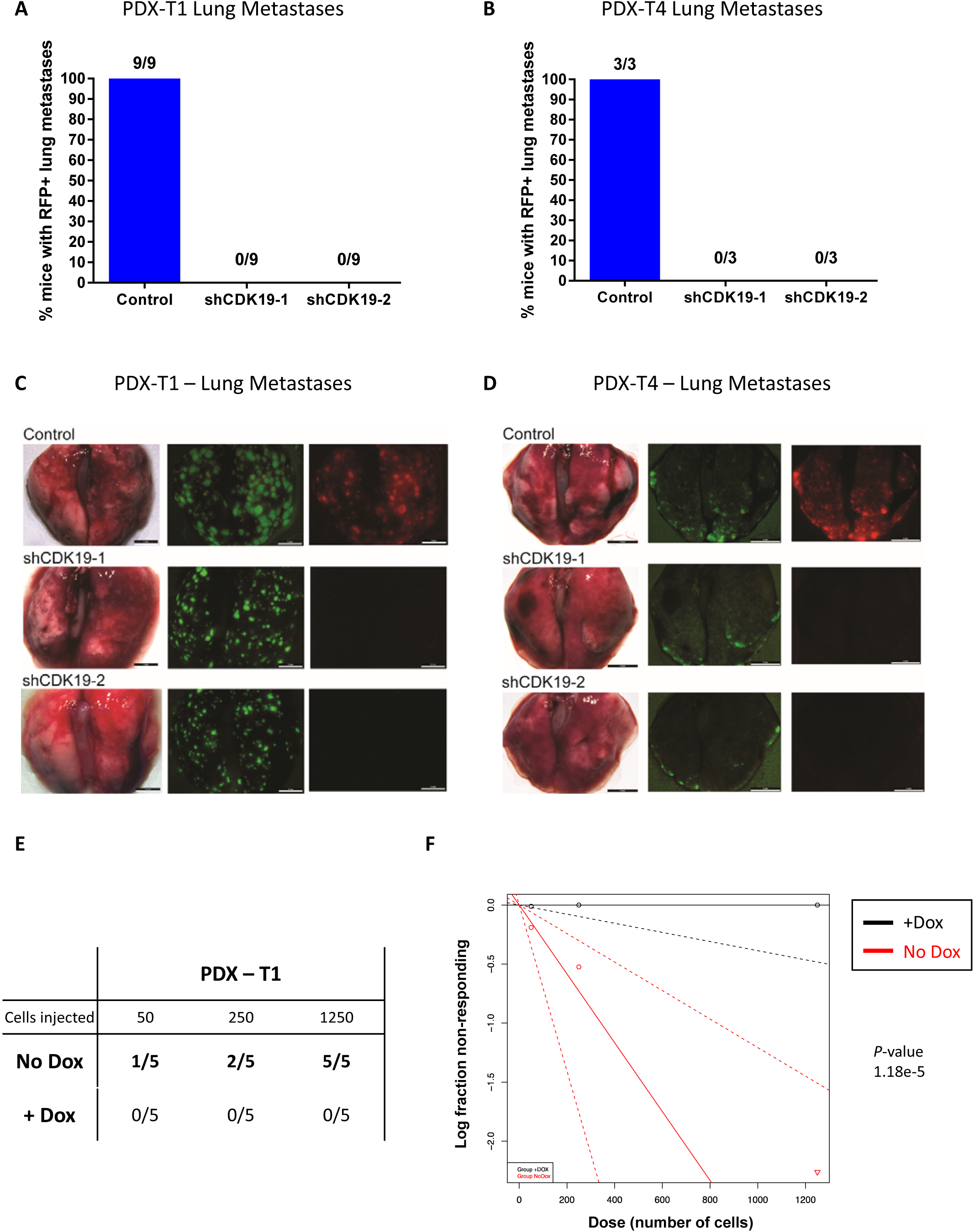
*CDK19* knockdown inhibits metastasis of TNBC cells and impairs tumor initiating capacity *in vivo*. (A and B) Percentage of mice with RFP positive lung metastases from mice bearing (A) PDX-T1 or (B) PDX-T4 primary tumor xenografts are shown. Number of mice with RFP positive lung metastases and total number of mice in each treatment group is shown as a fraction for each condition. PDX tumor cells were transduced with either control shRNA or *CDK19* targeting shRNA (shCDK19-1, shCDK19-2).
(C and D) Representative images of mouse lungs bearing (C) PDX-T1 or (D) PDX-T4 metastases. Lungs from mice with PDXs transduced with control shRNA (top row), shCDK19-1 (middle row) or shCDK19-2 (bottom row) are shown. Brightfield images (left column) show gross lung morphology, FITC images (middle column) identify metastatic tumor cells labeled with GFP (green) and Texas-Red images (right column) identify shRNA-transduced metastatic cells labeled with RFP (red). Scale bars = 5 mm.
(E) *CDK19* knockdown effectively prevents the growth of xenograft tumors in a limiting dilution assay. inducCDK19KD-PDX-T1 cells were injected into the mammary fat pads of NSG mice at 50, 250 and 1250 cells. Mice in the doxycycline group were fed a doxycycline containing rodent feed to induce *CDK19* shRNA, while mice in the control group were fed a normal rodent diet. Tumors were detected by palpation of tumors. The number of tumors that formed and the number of injections that were performed are indicated for each population. Populations and injections where tumors formed are bolded (5 mice for each condition tested).
(F) ELDA^21^ analysis of the data from (E) to determine tumor initiating frequencies in the doxycycline and control groups. *P*-value was determined by the ELDA software.

### CDK19 is important in maintaining tumor initiating capacity

Given that *CDK19* knockdown inhibited growth in two independent assays commonly used to assess tumorigenicity (PDX growth *in vivo* and organoid colony formation *in vitro*)^20^, and genes critical for tumor initiation are frequently enriched in specific sub-populations of cancer cells, we hypothesized that the tumor initiating cells (TICs) within the tumor might be sensitive to CDK19 inhibition. We sought to assess this with a limiting dilution assay (LDA) *in vivo* using inducCDK19KD-PDX-T1 cells where *CDK19* shRNA expression is activated exogenously with doxycycline food (Supplementary Fig. 3D). By comparing the *in vivo* transplantation of inducCDK19KD-PDX-T1 cells in the presence of doxycycline (+Dox) with the transplantation of inducCDK19KD-PDX-T1 cells without doxycycline (No Dox), we find that *CDK19* knockdown eliminated tumor formation regardless of the number of cells transplanted (Fig. 3E). Using Extreme Limiting Dilution Analysis software (ELDA)^21^, we determined that the tumor initiating frequencies significantly decreased from 1 in 342 cells (95%CI: 1 in 828 to 1 in 142) in the control (No Dox) group to 1 in ^∞^ cells (95%CI: 1 in ^∞^ to 1 in 2587) in the *CDK19* knockdown (+Dox) group (Fig. 3F). As *CDK19* knockdown significantly decreases tumor initiating frequency *in vivo* and reduces organoid colony formation *in vitro*, these findings suggested that CDK19 is important in maintaining the tumor initiating capacity of TNBC tumor cells from these PDXs.

### *CDK19* expressing cells also co-express tumor initiating cell-enriching surface marker genes

Previously, EpCAM and CD49f were utilized to isolate cell sub-populations in normal breast tissue^22^ and this approach was subsequently also applied to breast cancers^23^. With this strategy, EpCAM and CD49f have proven useful as markers to identify the EpCAM^med/high^ and CD49f^hi^ cells that enrich for TICs in TNBCs^24^. Using single cell qPCR analysis of the primary patient sample from which PDX-T1 was derived, we find that a significant portion (80%) of *CDK19* expressing cells also co-express both *EpCAM* and *CD49f* (Supplementary Fig. 4A). Thus, at the single cell level, this suggests a correlation between *CDK19* expression and the expression of markers that enrich for TICs.

### CD10 enhances isolation of tumor initiating cells

Despite the use of EpCAM and CD49f to identify TICs, we^24,25^ and others^23^ have found it difficult in many TNBC PDXs to use these markers to isolate cells into distinct sub-populations (Fig. 4A, left). Seeking to improve on this, we utilized the basal cell marker, CD10^26^ as an additional marker to FACS-sort breast cancer PDX’s to better isolate TICs. We discovered that CD10 and EpCAM separate TNBC PDX cells into three distinct sub-populations, EpCAM^med/high^/CD10^−/low^, EPCAM^low/med^/CD10^low/+^ and EpCAM^-^/CD10^−^ (Fig. 4A, right). Notably, the EpCAM^med/high^/CD10^−/low^ population corresponds well with the previously described EpCAM^med/high^/CD49f^high^ TIC enriched sub-population (Fig. 4B). To test the tumor initiating capacity of the three EpCAM/CD10 separated sub-populations, we performed organoid colony formation assays *in vitro* and transplantation limiting dilution assays (LDA) *in vivo*.

**Figure 4.**
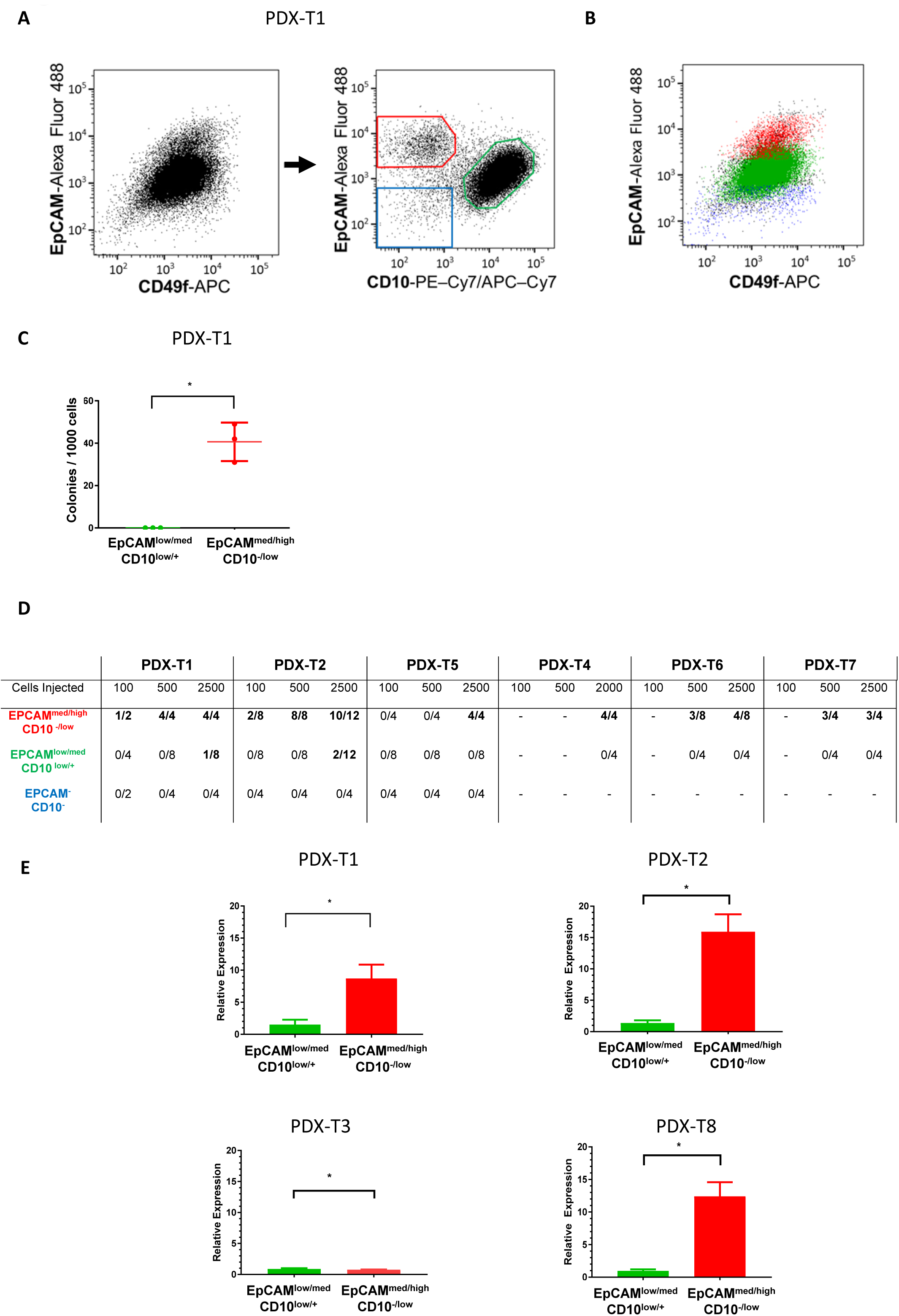
EpCAM and CD10 enables the isolation of a distinct tumor-initiating cell sub-population that overexpresses *CDK19*. (A) Representative flow cytometry analyses of a TNBC (PDX-T1) using EpCAM and CD49f (left) or EpCAM and CD10 (right) as cell surface markers. The large inseparable cell population (left) seen using EpCAM and CD49f, becomes three distinct sub-populations using EpCAM and CD10 (right): EpCAM^med/high^/CD10^−/low^ (red gate), EPCAM^low/med^/CD10^low/+^ (green gate) and EpCAM-/CD10-(blue gate).
(B) Flow cytometry analyses of TNBC (PDX-T1) using EpCAM and CD49f and the overlap of the EpCAM^med/high^/CD10^−/low^ (red), EPCAM^low/med^/CD10^low/+^ (green) and EpCAM-/CD10-(blue) sub-populations. Coloring as in (A).
(C) Comparing the organoid colony forming capabilities of the EpCAM^med/high^/CD10^−/low^ and EPCAM^low/med^/CD10^low/+^ cell sub-populations. The EpCAM^med/high^/CD10^−/low^ cells (red) formed significantly more organoid colonies than the EPCAM^low/med^/CD10^low/+^ cells (green), \**P* < 0.05 (unpaired t-test) (mean ± s.d., n = 3, experiment performed twice).
(D) PDX tumor cells were isolated by flow cytometry based on the expression of EpCAM and CD10 (as in Fig. 4A, right) and assayed for their tumor forming ability after injection into the mammary fat pads of NSG mice at 100, 500 and 2000 or 2500 cells. The number of tumors formed and the number of injections performed are indicated for each population. Populations and injections where tumors formed are bolded.
(E) *CDK19* expression is higher in the EpCAM^med/high^/CD10^−/low^ cells compared to the EPCAM^low/med^/CD10^low/+^ cells in PDX-T1, PDX-T2 and PDX-T8. Relative expression of *CDK19* in the EPCAM^low/med^/CD10^low/+^ (green) and the EpCAM^med/high^/CD10^−/low^ (red) cells as determined by RT-qPCR. Gene expression in each condition is normalized to beta-actin as a housekeeping gene. Relative expression of *CDK19* is normalized to the mean expression of *CDK19* in the EPCAM^low/med^/CD10^low/+^ cells. \**P* < 0.05 (unpaired t-test) (mean + s.d., n = 2 to 6. All experiments performed at least twice).

In *in vitro* organoid colony forming assays, the EpCAM^med/high^/CD10^−/low^ cells formed significantly more organoid colonies than the EpCAM^low/med^CD10^low/+^ cells (Fig. 4C). In transplantation assays performed in NSG mice, injection of EpCAM^med/high^/CD10^−/low^ cells from six TNBC PDXs (Supplementary Table 1) consistently formed tumors (Fig. 4D). In the case of PDX-T1 and PDX-T2, tumors formed with the transplant of as little as 100 cells. In contrast, transplant of EPCAM^low/med^/CD10^low/+^ cells did not form tumors in the majority of PDX tumors (PDX-T4, PDX-T5, PDX-T6, PDX-T7). In cases where tumors formed in the EPCAM^low/med^/CD10^low/+^ sub-population (PDX-T1 and PDX-T2), this only occurred when transplanting the highest cell numbers (i.e. 2500 cells). Furthermore, no tumors formed from the transplant of EpCAM^-^/CD10^−^ cells from *any* PDX. In secondary transplants of both PDX-T1 and PDX-T5, tumors only formed with the transplant of EpCAM^med/high^/CD10^−/low^ cells and not in the transplant of combined EPCAM^low/med^/CD10^low/+^ and EpCAM^-^/CD10^−^ cells (Supplementary Fig. 5). Thus, our data shows that TICs are enriched in the EpCAM^med/high^/CD10^−/low^ sub-population of all TNBC PDX breast tumors that we examined.

### *CDK19* expression is higher in the EpCAM^med/high^/CD10^−/low^ tumor initiating cell enriched tumor subpopulation

Given the overlap between *CDK19* expression and *EpCAM*/*CD49f* co-expression seen in our single cell data, we wanted to assess whether *CDK19* would also be expressed more highly in the tumorigenic EpCAM^med/high^/CD10^−/low^ cells compared to the less tumorigenic EPCAM^low/med^/CD10^low/+^ cells. In PDX-T1, PDX-T2 and PDX-T8, *CDK19* was expressed more highly in the more tumorigenic EpCAM^med/high^/CD10^−/low^ cells compared to the less tumorigenic EPCAM^low/med^/CD10^low/+^ cells (Fig. 4E). Thus, while *CDK19* was expressed in all the PDX tumors we examined, it was expressed at higher levels in the more tumorigenic EpCAM^med/high^/CD10^−/low^ sub-population in three of the four tumors that we examined.

### CDK19 uniquely regulates a subset of genes and pathways distinct from CDK8

Having identified a functional role for CDK19 in TNBC growth, we turned our attention to identifying potential genes and pathways regulated by CDK19. Despite early assumptions that CDK8 and CDK19 were redundant, gene knockdown studies in cell lines of other cancers^8,9^ have shown that CDK19 and CDK8 can regulate different genes. However, it is unclear whether this would also be the case in TNBC. TCGA data offered some evidence that the expression of *CDK19* and *CDK8* is independent in individual TNBC tumors. Patient tumors with *CDK19* CNA/mRNA upregulation are mutually exclusive from patient tumors with *CDK8* CNA/mRNA upregulation (Supplementary Fig. 4B). At the cellular level, our single cell qPCR data (Supplementary Fig. 4A) provided evidence that CDK8 and CDK19 are likely to have both shared and unique functions because only 49% of *CDK19* expressing cells also express *CDK8*. Through targeted knockdown of *CDK19* or *CDK8* and by thorough examination of the resulting global gene expression differences, we sought to investigate in TNBC whether CDK19 and CDK8 have redundant or biologically distinct functions.

We performed our investigations in MDA-MB231 cells where we knocked down *CDK19* or *CDK8* with constitutively expressed shRNA and examined the respective gene expression changes relative to control. Overall, *CDK19* knockdown affected 3909 genes and *CDK8* knockdown affected 4233 genes and both CDK19 and CDK8 positively and negatively regulated genes (Fig. 5A). However, only 12% of upregulated and 5% of downregulated genes in the *CDK19* knockdown experiment were also affected by *CDK8* knockdown. This suggested that although CDK19 has been considered redundant to CDK8, CDK19 and CDK8 largely regulate distinct genes (Fig. 5A). Gene Set Enrichment Analysis (GSEA)^27^ of the *CDK19* and *CDK8* knockdown genes allowed us to identify enriched hallmark gene sets^28^ that represent key biological processes or pathways amongst the most upregulated or downregulated genes (Fig. 5B and Supplementary Fig. 6). Genes associated with known breast cancer-related pathways such as mitosis (E2F targets, G2M Checkpoint, Mitotic Spindle), PI3K-AKT-MTOR signaling, MYC pathways (Myc Targets v1), glycolysis, apoptosis and oxidative phosphorylation were changed in the same direction by *CDK19* and *CDK8* knockdowns (Fig. 5B, violet, middle overlap region). These pathways appeared to be regulated by both CDK19 and CDK8. A number of pathways were only enriched in the genes that uniquely changed due to *CDK19* knockdown (Fig. 5B, left, blue region) or CDK8 knockdown (Fig. 5B, right, red region). Thus, while CDK19 and CDK8 regulate some of the same pathways, they each uniquely regulate their own set of pathways as well.

**Figure 5.**
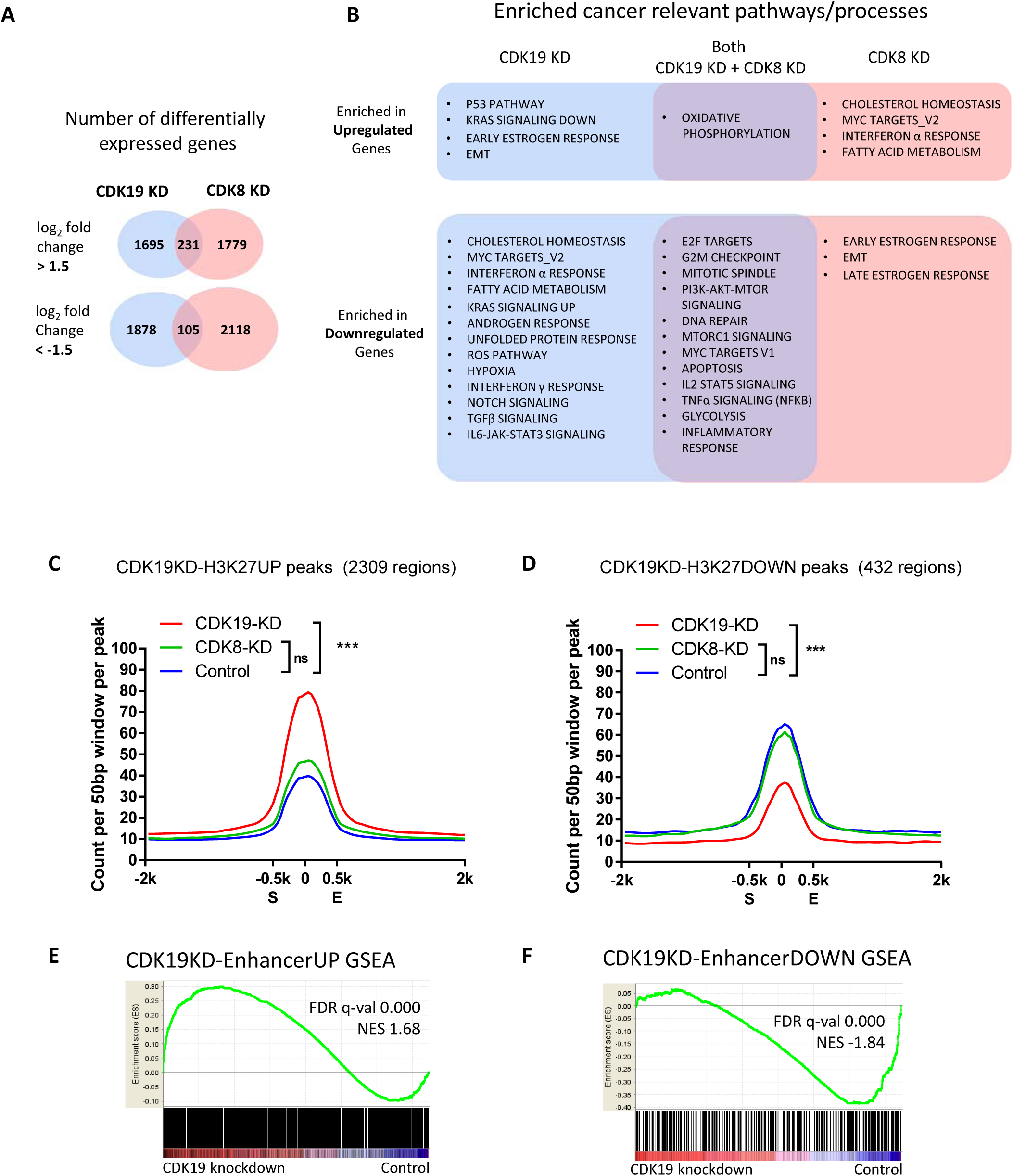
The effect of *CDK19* knockdown on gene expression, cancer relevant pathways and H3K27Ac. (A) Venn diagrams showing the number of genes upregulated *vs* control (upper diagram) and downregulated *vs* control (lower diagram) by *CDK19* knockdown (blue ovals), *CDK8* knockdown (red ovals) or by both *CDK19* knockdown and *CDK8* knockdown (overlap region).
(B) Venn diagram of cancer relevant pathways found to be enriched in the genes upregulated (upper diagram) or downregulated (lower diagram) by *CDK19* knockdown (blue region), *CDK8* knockdown (red region) or by both *CDK19* and *CDK8* knockdowns (overlap region) as determined by GSEA. The full list of enriched pathways and the associated statistics are shown in Supplementary Fig. 6.
(C and D) CHIP-Seq signals across the CDK19KD-H3K27AcUP and CDK19KD-H3K27AcDOWN regions are significantly different in the *CDK19* knockdown samples compared to control. Aggregate plots of normalized H3K27Ac CHIP-Seq signal across (C) CDK19KD-H3K27UP regions or (D) CDK19KD-H3K27DOWN regions in the *CDK19* knockdown (blue line), *CDK8* knockdown (green line) and control (red line) samples, \*\*\**P* < 0.001; ns is *P* > 0.05 (all samples n = 3, experiments performed three times). H3K27Ac CHIP-Seq signals of the CDK19KD-H3K27UP or CDK19KD-H3K27DOWN regions are normalized to 1-Kb and centered on the middle of those regions (S and E denote start and end, respectively). Signals of the flanking 2-Kb regions are also shown. To compare relative signal changes, the total signal of each biological replicate was determined by summing the signals of each 50-base window 1-Kb around the center of each region. *P*-values between total CHIP-Seq signals of each sample were determined by unpaired t-test.
(E and F) GSEA of (E) CDK19KD-EnhancerUP or (F) CDK19KD-EnhancerDOWN genes using averaged *CDK19* knockdown versus Control expression data.

### H3K27Ac modulation from *CDK19* knockdown correlates with gene expression at putative gene enhancers

Recent studies have highlighted the role of CDK19 and CDK8^5^, as well as other transcriptional CDKs (CDK7^29^, CDK12/CDK13^30^), in regulating the transcription of critical oncogenic genes by acting at large clusters of enhancers (also called ‘super-enhancers’) that are marked by histone 3 lysine 27 acetylation (H3K27Ac)^31^. The exact mechanism for this gene regulation is unclear, but is believed to occur in part through interactions of the CKM with core Mediator to regulate RNAPII-Mediator interactions^8^ and in part by phosphorylating serine residues in the C-terminal domain of RNAPII^1^. Given the propensity of transcriptional CDKs to function at enhancers, we wanted to investigate genome-wide whether CDK19 and CDK8 can regulate histone acetylation as a mechanism to control gene expression. While enhancer modification through other signaling pathways have been identified^32,33^, this mechanism of gene control has not yet been reported for the CDKs.

To explore the role of CDK19 in epigenetic regulation, chromatin immunoprecipitation and sequencing (CHIP-Seq) for the H3K27Ac modification was performed on MDA-MB231 cells under three different conditions: *CDK19* knockdown, *CDK8* knockdown and control (empty vector transduction). Genome-wide analysis of *all* H3K27Ac modified regions showed that both *CDK19* knockdown and *CDK8* knockdown had similar global H3K27Ac levels compared to control (Supplementary Fig. 7A). Through comparative analysis of H3K27Ac levels in the *CDK19* knockdown compared to the control, we identified 3034 regions with increased H3K27Ac signal (All-H3K27UP) and 502 regions with decreased H3K27Ac signal (All-H3K27DOWN). By excluding regions that also differed between *CDK8* knockdown and control, we identified 2309 regions unique to *CDK19* knockdown with increased H3K27Ac signal (CDK19KD-H3K27UP) and 432 regions with decreased H3K27Ac signal (CDK19KD-H3K27DOWN). The specificity of these regions for CDK19 was investigated by comparing the H3K27Ac levels at these regions in *CDK19* knockdown, *CDK8* knockdown and control samples. Compared to control, enrichment of H3K27Ac levels across the CDK19KD-H3K27UP regions (Fig. 5C) and depletion of H3K27Ac levels across the CDK19KD-H3K27DOWN regions (Fig. 5D) were significant only for *CDK19* knockdown and not for *CDK8* knockdown. Thus, CDK19KD-H3K27UP and CDK19KD-H3K27DOWN define regions where the H3K27Ac signal is more specific for, and most sensitive to, knockdown of *CDK19* compared to knockdown of *CDK8*.

We next assessed whether increases or decreases in H3K27Ac levels as a result of *CDK19* knockdown corresponded to changes in gene expression. For this, the previously defined All-H3K27UP and All-H3K27DOWN regions were annotated by proximity to the nearest gene to establish two gene sets: CDK19KD-EnhancerUP (1593 genes) and CDK19KD-EnhancerDOWN (341 genes) for further analyses (Supplementary Table 2). Enrichment analysis of these gene sets with our *CDK19* knockdown gene expression data indicated that genes most upregulated by *CDK19* knockdown were enriched for the CDK19KD-EnhancerUP genes (NES 1.68, FDR q-value = 0.000) (Fig. 5E), while genes most downregulated by *CDK19* knockdown were enriched for the CDK19KD-EnhancerDOWN genes (NES −1.84, FDR q-value = 0.000) (Fig. 5F). An examination of sample genes shows that increases (Supplementary Fig. 7B and 7C) or decreases (Supplementary Fig. 7D and 7E) in H3K27Ac enhancer signals correlate with corresponding changes in gene expression (Supplementary Fig. 7F). Thus, as a result of *CDK19* knockdown, perturbations to the H3K27Ac signal at the putative enhancer elements of genes correlated well and in the expected direction with changes in gene expression.

### *CDK19* knockdown in pre-established tumors suppressed the growth of PDX tumors and prolonged survival

To explore translational applications of CDK19 targeting, we modelled treatment of patient tumors by investigating the effect of *CDK19* knockdown on the growth of pre-established organoids *in vitro* and pre-established PDX tumors *in vivo*. *In vitro*, using inducCDK19KD-PDX-T1 cells to pre-establish organoid colonies (Fig. 6A, left), the induction of *CDK19* shRNA in the doxycycline group significantly decreased the number of organoids compared to the Control group (without doxycycline) (Fig. 6A, right). *In vivo*, in NSG mice in which inducCDK19KD-PDX-T1 or inducCDK19KD-PDX-T3 tumors were pre-established, introduction of doxycycline into the mouse feed to induce *CDK19* knockdown significantly impaired the growth of these tumors (Fig. 6B and 6C). *CDK19* shRNA induced tumors were ultimately 82% smaller in inducCDK19KD-PDX-T1 tumors (Fig. 6B) and 38% smaller in inducCDK19KD-PDX-T3 tumors (Fig. 6C) when compared to control tumors. In both the inducCDK19KD-PDX-T1 and inducCDK19KD-PDX-T3 experiments, mouse total body weights were not significantly different between the treatment and control groups (Supplementary Fig. 8A and 8B).

**Figure 6.**
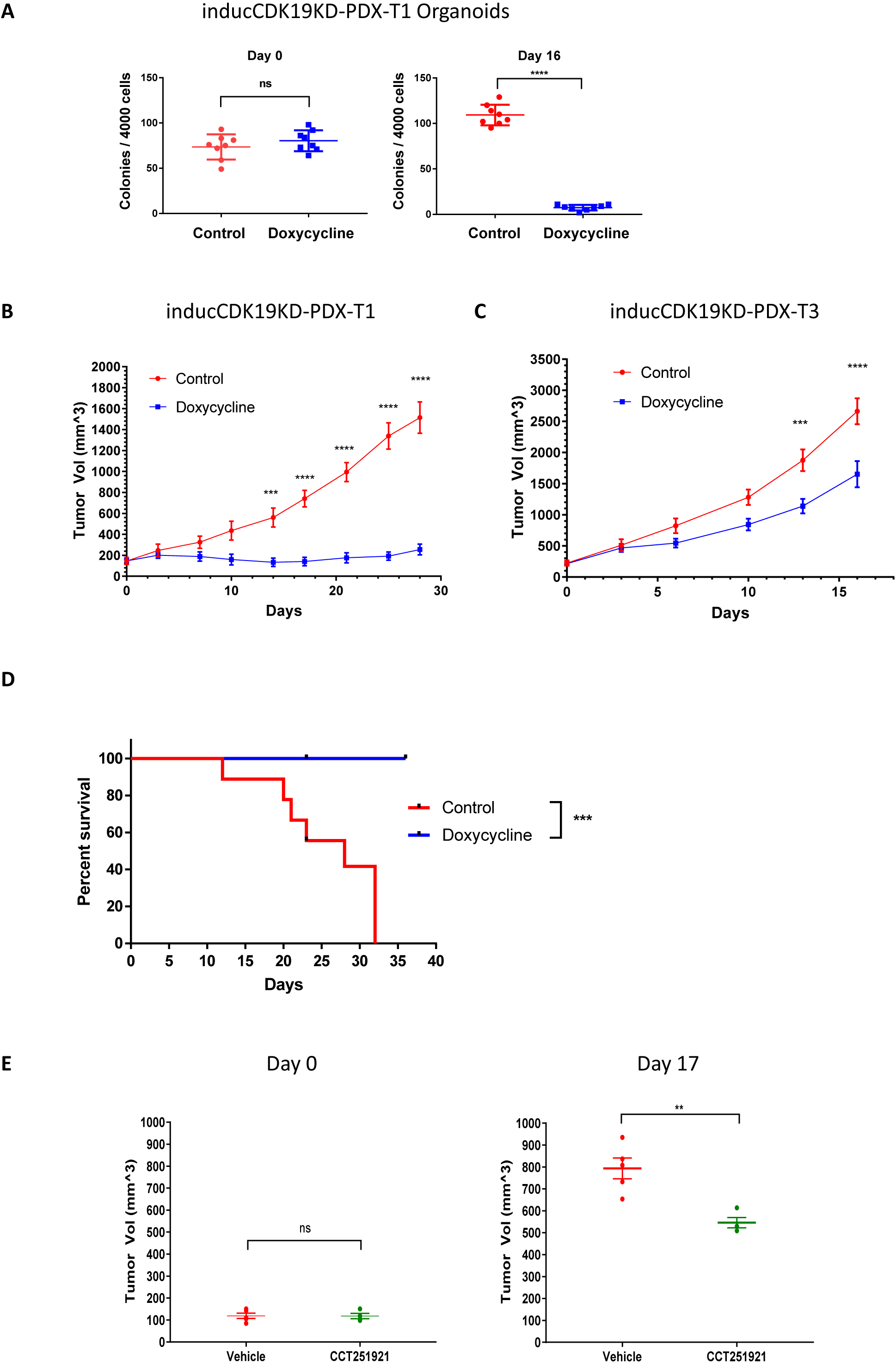
Genetic knockdown of *CDK19* or pharmacologic inhibition of CDK19 kinase activity can impair growth of pre-established tumors. (A) In inducCDK19KD-PDX-T1 cells, induction of *CDK19* shRNA by addition of doxycycline significantly decreased the number of organoid colonies in the doxycycline treatment group (blue) compared to control (red). Number of organoid colonies at Day 0 (left) and Day 16 (right) after initiating doxycycline treatment is shown, \*\*\*\**P* < 0.0001; ns is *P* > 0.05 (mean ± s.d., n = 6, experiment performed twice, *P*-values determined by unpaired t-test).
(B and C) Induction of *CDK19* shRNA in pre-established tumors impaired tumor growth. The growth of pre-established tumors in the doxycycline fed NSG mice (blue) and control NSG mice (red) are shown for (B) inducCDK19KD-PDX-T1, \*\*\**P* < 0.001; \*\*\*\**P* < 0.0001 (mean ± s.e.m, total of 10 mice for each condition tested) and (C) inducCDK19KD-PDX-T3, \*\*\**P* < 0.001; \*\*\*\**P* < 0.0001 (mean ± s.e.m, total of 10 mice for each condition tested). All *P*-values determined by 2-way ANOVA.
(D) Kaplan-Meier survival curves for mice engrafted with inducCDK19KD-PDX-T1 xenografts. NSG Mice with pre-established tumors (as described in Figure 6B) were fed normal food (Red Line) or doxycycline food to induce CDK19 knockdown (Blue Line). Mice tumor volumes were followed with measurements every three to four days. Mice were sacrificed when the longest diameter of their tumor exceeded 17 mm, ***P < 0.001 (10 mice for each condition tested, log-rank (Mantel-Cox) test used to determine P-values).
(E) Treatment of mice with CCT251921 by daily oral gavage significantly impaired the growth of pre-established PDX-T1 xenograft tumors. NSG mice with pre-established PDX-T1 xenograft tumors were treated with daily oral gavage of CCT251921 (green) or vehicle (red). Tumor volumes at day 0 (beginning) and day 17 (end of the experiment) are shown, ns is *P* > 0.05, \*\**P* < 0.01; (mean ± s.e.m., 4 to 5 mice for each condition tested, *P*-values determined by unpaired t-test).

To model clinical outcomes we also performed survival studies in mice (based on time for tumors to grow to a pre-determined size before mandatory euthanization). We found that mouse survival was significantly longer in inducCDK19KD-PDX-T1 mice whose PDX tumors had *CDK19* knocked down compared to control mice (Fig. 6D). In summary, these experiments showed that *CDK19* can significantly decrease the growth of pre-established tumors and can prolong survival in mice.

### Chemical Inhibition of CDK19 and CDK8 in pre-established tumors suppressed the growth of PDX tumors

To model the use of a CDK19 targeted therapies clinically, we utilized CCT251921^34^ (Supplementary Fig. 8C), an orally bioavailable inhibitor of both CDK19 and CDK8. PDX-T1 tumors were pre-established in NSG mice (Fig. 6E, left) before starting daily oral administration (30mg/kg) of CCT251921 or vehicle. Treatment with CCT251921 over 17 days resulted in significant reductions in final tumor volumes compared to tumors in mice treated with vehicle (Fig. 6E). Final volumes of tumors in CCT251921 treated mice were over 30% smaller than the tumors of vehicle treated mice (Fig. 8E). The weights of mice in both the CCT251921 and vehicle experiments decreased slightly, but were not significantly different between conditions at the end of the experiment (Supplementary Fig. 8D). It is well known that different biological outcomes can arise from gene knockdown versus chemical inhibition^35^. We show here in pre-established tumors that chemical inhibition of CDK19 kinase activity can recapitulate some, but not all, of the effects of total *CDK19* loss shown in our knockdown studies.

## Discussion

Here, through a combination of *in vitro* studies using PDX organoids and *in vivo* studies using PDX tumors in NSG mice, we comprehensively validated the inhibitory effects of *CDK19* knockdown in TNBC. Notably, *CDK19* knockdown appeared to have the greatest effect on tumor initiation as observed through our finding of reduced organoid colony formation *in vitro* and reduced tumor transplantation *in vivo*. Using CD10 and EpCAM as a novel surface marker combination to isolate distinct TNBC sub-populations, we found that in many tumors the TIC-enriched EpCAM^med/high^/CD10^−/low^ sub-population expresses *CDK19* more highly than other sub-populations. This suggests that TICs are likely more sensitive to CDK19 loss and putatively the cellular targets of *CDK19* knockdown. In addition to CDK19’s role in tumor initiation, our studies highlight a role for CDK19 in tumor proliferation because *CDK19* knockdown also inhibited the growth of pre-established PDXs. Finally, we also found that *CDK19* knockdown inhibited tumor metastases. This did not correlate with primary tumor volume (Supplementary Fig. 3E) or time since appearance of primary tumor (Supplementary Fig. 3F), suggesting that it is related to impairment of tumor metastatic potential. Thus, in TNBC, CDK19 appears to play a role in multiple aspects of cancer growth including tumor initiation, proliferation and metastases. This does not exclude the possibility that CDK19 is important in other breast cancer subtypes as well. In fact, the significant correlation between *CDK19* expression and relapse free survival in ER+ breast cancer patients (Supplementary Fig. 2E) and the inhibitory effect of CDK19/8 inhibitors in ER+ cell lines^36^ suggests a possible role for CDK19 in the ER+ subtype as well.

Our gene expression studies in TNBC showed that like CDK8^1,5^, CDK19 can both positively and negatively regulate genes and pathways. While CDK19 regulates some pathways in common with CDK8, it also uniquely regulates its own distinct set of genes and pathways. This shows that CDK19 does not simply serve a redundant role to CDK8 as once thought. The finding that CDK19 regulates unique genes and pathways parallels our data (Fig. 5B) and prior data showing that CDK8 can also mediate its own unique pathways, such as the HIF1A pathway^8^. Molecular studies showing that CDK8 and CDK19 bind mutually exclusively to core Mediator^8^ can perhaps explain some of the distinctiveness of the pathways that they regulate. On the other hand, the overlap of pathways regulated by both CDK19 and CDK8 may derive partially from the sharing of common CKM components (Cyclin C, Med12, Med12L, Med13 and Med13L) to form different CKM variants^3^.

Unsurprisingly, a review of the pathways uniquely regulated by CDK19 (Fig. 5B) in TNBC revealed a number of pathways such as cholesterol homeostasis, P53 signaling, mitosis and NFκB that have previously been associated with CDK19 in other cancer cell types^37,38^. Notably, many CDK19 unique pathways that we uncovered such as P53 signaling, KRAS signaling and others are important to the biology of TNBC^39^ and are actively being pursued in TNBC clinical trials^40^. CDK19 might therefore be a potential regulator of multiple pathways that are important in driving TNBC growth.

Complementing studies showing that CDK8 and other CDKs specifically function at ‘super enhancers’ marked by H3K27Ac^5,29,30^, our CDK19 knockdown CHIP-Seq studies showed a potential role for CDK19 in regulating H3K27Ac levels. Genome-wide, H3K27Ac modulation from *CDK19* knockdown impacted gene expression as seen in the strong correlations noted from enrichment analysis (Fig. 5E and 5F). Further exploration of the mechanisms by which CDK19 modulates H3K27Ac levels and the functional consequences of these modifications on signaling pathways would be interesting. Although we show here the first example of CDK19 mediating H3K7Ac changes, previous studies have shown other roles for CDK19 in epigenetic regulation. CDK19 (and CDK8) can interact with PRMT5 to regulate methylation of histone H4 arginine 3 in gene promoters to repress gene expression^41^. Thus, in addition to regulating transcription through more direct CDK19-RNA polymerase II mechanisms, epigenetics represents another means by which CDK19 can modulate gene expression.

As a clinical target for TNBC, CDK19 appears to have significant potential on many fronts. The multitude of clinically relevant TNBC pathways regulated by CDK19 suggests that targeting CDK19 could provide the opportunity to modulate multiple pathways simultaneously. This approach could overcome the resistance to single agent therapy commonly seen in TNBC^40^ and also potentially enable the targeting of ‘undruggable’ pathways such as those involving P53 or MYC. The potential to inhibit CDK19, a transcriptional co-factor, to target TICs is equally exciting. Current strategies inhibiting transcriptional co-factors utilize targets such as other CDKs or BRD4, while strategies targeting TICs have focused on inhibiting self-renewal pathways such as Hedgehog, Wnt/β-catenin and Notch. Unfortunately, both these therapeutic approaches are often limited by toxicity to normal cells. This can be attributed to the ubiquitous expression of transcriptional co-factors in normal tissues and the importance of self-renewal pathways in normal stem cells. BRD4 inhibition, for example, resulted in a disruption of tissue homeostasis in multiple organs in mice^42^. Similarly, due to the challenge of narrow therapeutic indices, Hedgehog, Notch and Wnt pathway inhibitors have all had limited clinical success thus far^43^.

Our data showing that *CDK19* knockdown does not affect untransformed mammary epithelial cells (Fig. 1D), and the known biology of CDK19 suggests potential advantages of CDK19 as a therapeutic target. Compared to other ubiquitous transcriptional co-factors such as CDK8, CDK9 and BRD4, CDK19 has more limited tissue distribution^9^, potentially limiting the toxicity from systemic CDK19 inhibition. Supporting this notion is the observation that *CDK8*, *CDK9* and *BRD4* knockouts are lethal^11,12,44^ whereas *CDK19* knockout mice are viable^12^. In addition, the limited expression of CDK19 in sensitive tissues such as colon (Supplementary Fig. 1B) could broaden the therapeutic window for its activity.

To explore the translational potential of CDK19 inhibitors for human therapy, we investigated the effect of an oral CDK19/CDK8 dual inhibitor on the growth of pre-established PDXs in mice. This showed a modest reduction of tumor growth in treated mice (Fig. 6E). However, these results were confounded by the inhibition of both CDK19 and CDK8 kinases by the dual inhibitor. Ideally, we would utilize a CDK19-specific inhibitor, but such an inhibitor has yet to be reported. Most CDK19 kinase inhibitors reported to-date including CCT251921, Senexin A and Cortistatin A have been equipotent on both CDK19 and CDK8^5,18,34^. Promisingly, the recent report of a CDK8-specific inhibitor^45^ suggests that sufficient differences might exist between the CDK19 and CDK8 kinase domains for CDK19-specific inhibition to be feasible.

We compared *CDK19* knockdown to dual CDK19/CDK8 kinase inhibition to gain some insight into whether CDK19 protein loss and CDK19 kinase inhibition have similar effects. The modest effect of our dual inhibitor on tumor growth (Fig. 6E) compared to the more pronounced effect of genetic knockdown (Fig. 6B) suggests that kinase inhibition does not fully re-capitulate the effects of protein loss. This discrepancy is consistent with studies showing that the outcomes from dual CDK19/CDK8 genetic knockdown and dual kinase chemical inhibition can differ^35^. Thus, small molecule strategies targeting proteins for degradation^46^ might be necessary to fully recapitulate the effects seen from genetic knockdown of *CDK19*.

In conclusion, the data presented here and the known biology of CDK19 suggests that targeting CDK19 holds significant promise for TNBC treatment. However, many difficult challenges regarding CDK19 specificity and inhibition strategies need to be resolved before targeting CDK19 can become a reality in the clinical setting.

## Methods and Materials

### Chemical Reagents

The following are the chemical names for the compounds used in this study. **CCT152921** is 4-[(2-Phenylethyl)amino]-6-quinazolinecarbonitrile (NIH NCAT). Compound was re-suspended in Vehicle (PBS + 0.5% Methocel (w/v) + 0.25% Tween 20 (v/v)) to a concentration of 3mg/mL and mice were dosed at 30mg/kg^34^. CCT251921 or vehicle was administered via daily oral gavage.

### shRNA expression lentiviral plasmids

Pairs of complementary ssDNA oligonucleotides containing the sense target sequence, a 15-mer loop sequence (5′-GTTAATATTCATAGC-3′) and the reverse complement of the sense sequence were synthesized (Elim Biopharmaceuticals). The oligos were annealed in 50 μM annealing buffer (10 mM Tris-HCl pH 8.0, 50 mM NaCl, 1mM EDTA). The double-stranded DNA oligo templates were subsequently cloned into the pRSI12-U6-(sh)-HTS4-UbiC-TagRFP-2A-Puro shRNA expression vector (Cellecta) digested with *BbsI* for constitutively active shRNA vector constructs and pRSITUR-U6Tet-(sh)-UbiC-TetRep-2A-TagRFP digested with *BbsI* for inducible shRNA vector constructs. The shRNA sequences used in this study to construct knockdown vectors were (sense and anti-sense strands are bolded): 5′-**GCG AGA ATT GAA GTA CCT TAA** – GTT AAT ATT CAT AGC – **TTA GGG TGC TTC AAT TCT CGC**-3′ (shCDK19-1), 5′-**ACC AGC AAA TAT CCT AGT AAT** – GTT AAT ATT CAT AGC – **ATT ACT AGG ATA TTT GCT GGT**-3′ (shCDK19-2) and 5’-**GCA GGG TAA TAA CCA CAT TAA** – GTT AAT ATT CAT AGC – **TTA GTG TGG TTA TTG CCC TGC**-3’ (shCDK8-2). Only shCDK19-1 was used in the inducible constructs as the effects from using shCDK19-1 and shCDK19-2 appeared functionally similar in PDXs. The unmodified pRSI12-U6-(sh)-HTS4-UbiC-TagRFP-2A-Puro shRNA and pRSITUR-U6Tet-(sh)-UbiC-TetRep-2A-TagRFP expression vectors above were used as the ‘empty’ control shRNA. The pHIV-ZsGreen expression vector (Addgene) was used to produce GFP positive tumor cells.

### Cell Lines

MDA-MB231, MDA-MB468 and 293T cells were obtained from ATCC. HMEC cells were obtained from ThermoFisher Scientific. These cells were certified by the vendors to be mycoplasma free. None of the cell lines used are listed in the database of commonly misidentified cell lines maintained by ICLAC. Cell lines have not been authenticated but all cell lines used were passaged less than 10 times from when the original cells from the vendors were thawed. Cell lines have not been tested for mycoplasma contamination. MDA-MB231, MDA-MB468 and 293T cells were grown in DMEM (Invitrogen) supplemented with PSA (Life Technologies), 10% FBS (Hyclone), Glutamax (ThermoFisher Scientific) and sodium pyruvate (Life Technologies). HMEC cells were grown in HuMEC Ready Medium (ThermoFisher Scientific).

### Mice

Nod *scid* gamma (NSG) mice (NOD.Cg-Prkdc^scid^ IL2Rg^tm1Wjl^/SzJ) were purchased from the Jackson Laboratory. Mice used for PDX experiments were adult female mice between 8 – 10 weeks old. All the mice used in this study were maintained at the Stanford Animal Facility in accordance with a protocol approved by the Stanford University APLAC committee. Mice were maintained in-house under aseptic sterile conditions. Mice were administered autoclaved food and water. For PDX experiments utilizing doxycycline inducible constructs, mice were provided rodent feed containing 625mg Doxycycline hyclate/kg diet (Envigo) in place of their normal rodent diet.

### PDX Tumors and their pathological and clinical characteristics

For human samples, informed consent was obtained after the approval of protocols by the Institutional Review Boards of Stanford University and The City of Hope. See Supplementary Table 1 for a full description of all the PDX tumors used in this study.

### Single cell suspension of PDX tumor cells

Xenografts were mechanically chopped with a razor blade to approximately 1 mm pieces and then incubated at 37 degrees Celsius for 3 to 4 hours with collagenase and hyaluronidase (Stem Cell Technologies) in Advanced DMEM/F12 (Invitrogen) with 120 μg/ml penicillin, 100 μg/ml streptomycin, 0.25 μg/ml amphotericin-B (PSA) (Life Technologies). Cells were then treated with ACK lysis buffer (Gibco) to lyse red blood cells, followed by 5 mins of treatment with pre-warmed dispase (Stem Cell Technologies) plus DNAseI (Sigma) and filtered through a 40 μm nylon mesh filter. Cells were finally washed with flow cytometry buffer (HBBS, 2% FCS, PSA).

### Lentivirus Production

Lentivirus was produced with Packaging Plasmid Mix (Cellecta) and subcloned pRSI12 or pRSITUR shRNA expression plasmids using Lipofectamine 2000 (Thermofisher Scientific) in 293T cells per manufacturer’s instructions. Supernatants were collected at 48 h and 72 h, filtered with a 0.45 μm filter and precipitated with Lentivirus Precipitation Solution (Alstem LLC) per manufacturer’s instructions. Virus was resuspended in 1/100 original volume. Viral titers were determined by flow cytometry analyses of 293T cells infected with serial dilutions of concentrated virus.

### Lentivirus Infection

For *in vitro* cell line experiments, concentrated lentiviral supernatant (to achieve an MOI of 3) was mixed with cells at the time of seeding. Cells were monitored by visualization of RFP under fluorescence microscopy. All flow cytometry analyses were performed after at least 72 hours of infection.

For *in vivo* PDX tumor growth and organoid colony formation experiments, concentrated lentiviral supernatant (to achieve an MOI of 10) was mixed with single cell suspensions of PDX tumor cells in organoid media with 4 μg/ml of Polybrene (Sigma-Aldrich). Organoid media consisted of: Advanced DMEM/F12 (Invitrogen), 10% FBS (Hyclone), 2.5% growth factor-reduced Matrigel (BD), 10 ng/ml mouse EGF (R&D), 100 ng/ml Noggin (R&D), 250 ng/ml RSPO-I (R&D), 1× B27 (Invitrogen), 1× N2 (Invitrogen) and PSA (Life Technologies). Cells were then spinoculated by centrifuging at 15 degrees Celsius for 2 hours at 1200×*g*. Cells were resuspended by pipetting and left overnight in 48-well ultra-low attachment cell culture plates (Corning).

For organoid colony formation assays, cells were transferred the next day to matrigel. For *in vivo* PDX assays, approximately 75% of the cells were injected into NSG mice as described in the PDX tumor engraftment section. The remainder 25% of cells were plated on matrigel and grown in organoid media for 72 hours until the cells became RFP positive. At that point media was removed and exchanged for dispase and incubated for 2 – 3 h until the matrigel dissolved. Dissociated cells were resuspended in flow cytometry buffer and analyzed by flow cytometry to determine the ‘baseline’ RFP percentage for cells that were injected into the mice.

### Organoid Colony Formation Assay

Irradiated L1-Wnt3a feeder cells (generous gift of Dr. Roel Nusse) were mixed with growth factor reduced matrigel (BD Biosciences) and allowed to solidify at 37 degrees Celsius. Single cell suspensions of PDX tumor cells were transferred onto the solidified matrigel/feeder cell mix substrate and grown in organoid media. Cells were grown for approximately 2 weeks in a 37 degrees Celsius incubator with 5% CO2. 50% of media was exchanged with fresh media every 3 – 4 days. Colonies were counted under fluorescence microscopy. Only RFP positive colonies (which represent transduced cells) were counted. For experiments in which we induced expression of CDK19 shRNA, doxycycline hyclate was added to a final concentration of 100ng/mL into the media.

### Cell Viability Assay

For cell lines treated with chemical or infected with lentivirus, WST-1 Cell Proliferation Reagent (Roche) was added at 1:10 (v/v) final dilution to each well per manufacturer’s instructions. Cells were subsequently incubated at 37 degrees Celsius and 5% CO2. Between 1 – 4 hours after addition of reagent, plates were analyzed on a SpectraMax M3 Bioanalyzer (Molecular Devices). Absorbance for each well was measured at 450nm (signal wavelength) and 650nm (reference wavelength). Thus, the signal for each experimental sample was Absorbance_experimental_ (A_450nm_-A_650nm_). To correct for the effect of media, Absorbance_background_ (A_450nm_-A_650nm_) was obtained by measuring absorbance in a blank well. Thus, the background corrected signal for each sample A_corrected_ = Absorbance_experimental_ – Absorbance_background_. All A_corrected_ values for the knockdowns were normalized to the A_corrected_ value for the control sample to obtain a ‘Relative Viability’.

### Quantitative PCR RNA expression analyses

Cells were lysed with Trizol (Life Technologies) and RNA was extracted according to the manufacturer’s instruction. RNA was then treated with *DNAseI* to remove contaminating genomic DNA. RNA was reverse transcribed to cDNA using SuperScript III First Strand Synthesis kit (Life Technologies) according to the manufacturer’s instructions. TaqMan Gene Expression Master Mix (Applied Biosystems) and the following TaqMan Gene Expression Assays (Applied Biosystems) were used following manufacturer’s instructions: *ACTB*, Hs00357333_g1 and *CDK19*, Hs01039931_m1. Data was collected on a 7900HT Fast Real-Time PCR System (Applied Biosystems) and data analyzed with SDS 2.4 software (Applied Biosystems). Gene expression data in each sample was normalized against the expression of beta-actin.

### PDX tumor cell engraftment, limiting dilution assays and secondary transplantation assays

Single cell suspensions of PDX cells were resuspended in 50% (v/v) mixtures of normal matrigel (BD Biosciences) and flow cytometry buffer in a total volume of 50 – 100 μl. Using an insulin syringe, cells were injected subcutaneously into the nipple of female NSG mice at the fourth abdominal fat pad.

For limiting dilution assays and secondary transplantion assays, dissociated cells from PDX tumors were sorted with EpCAM and CD10 into the following sub-populations: EpCAM^med/high^/CD10^−/low^, EPCAM^low/med^/CD10^low/+^ and EpCAM^-^/CD10^−^ as shown in Fig. 4A. The specific number of cells injected into the mice were determined by flow cytometry and secondarily by manual counting with a hemocytometer.

### PDX tumor growth and mice total body weights

PDX tumors were detected by palpation. Tumor volumes were determined by measuring the length (l) and width (w) and calculating volumes using the ellipsoid formula 1/6 × l × w^2^ × π.

### Mouse PDX tumor and lung dissection

Xenograft tumors and mice lungs were surgically resected after the mice were euthanized. A 3 to 4 mm section was cut from each tumor and saved in ice cold PBS for imaging. The mice lungs and tumors were imaged on a M205FA Fluorescence Stereo Microscope (Leica) and images were captured with a DFC310FX camera (Leica).

### Flow cytometry to determine RFP percentage

Flow cytometry was performed with a 100 μm nozzle on a Flow Cytometry Aria II (BD Biosciences) with Diva software (BD Biosciences). Data analysis was performed using Flowjo software (Flowjo). For all experiments, side scatter and forward scatter profiles (area and width) were used to eliminate debris and cell doublets. Dead cells were eliminated by excluding 4′,6-diamidino-2-phenylindole (DAPI)-positive cells (Molecular Probes). For PDX tumor cells, they were gated for GFP positivity and then for RFP positivity. RFP percentage is the percentage of GFP positive cells that are also RFP positive. For each sample, we obtain the RFP fraction that is: the RFP % in the tumor divided by the baseline RFP % (see ‘Lentivirus infection’ section). RFP fraction for each sample is then normalized to the RFP fraction for the shRNA control sample which is set at 100% to obtain the ‘Normalized % RFP’.

### Flow Cytometry using EpCAM, CD10 and CD49f cell surface markers for analysis and cell sorting

Flow cytometry for analysis and cell sorting was performed as previously described^25^. Human antibodies used included: EpCAM–Alexa Fluor 488 (clone 9C4, Biolegend); 1 μg ml^−1^, CD49f–APC (clone GoH3, Biolegend); CD10 PE-Cy7/APC–Cy7 (clone H110a, Biolegend); 1 μg ml^−1^ and H-2Kd biotin/Pacific Blue (clone SF1-1.1, Biolegend); 1 μg ml^−1^.

### RNAi Dropout Viability Screen

GFP positive PDX-T1 tumors grown in NSG mice were dissected, processed to single cells and enriched with EpCAM as described previously. Analysis of cells at this point showed that they were approximately 98% - 100% GFP positive.

For the *in vitro* RNAi dropout viability screen, 60 million dissociated PDX-T1 cells were transduced with the DECIPHER 27K Pooled shRNA lentivirus library–Human Module 1 (Cellecta) at an MOI of 1 (based on 293T cell tittering) in the presence of polybrene and then spinoculated for 2 hours as described previously. The next day, half the cells were spun down and frozen as the *in vitro* baseline reference sample. The remainder of the cells were plated into twelve 150 mm dishes prepared with 12 ml matrigel containing irradiated L1-Wnt3a feeder cells at 250,000 cells/ml of matrigel. The cells were grown for 19 days with an exchange for fresh media every 3-4 days. On the final day, all the media was exchanged with dispase in order to dissolve the matrigel and to recover the cells. The cells from all the plates were pooled, washed and frozen as the *in vitro* organoid growth experimental sample.

For the *in vivo* RNAi dropout viability screen, 30 million dissociated PDX-T1 cells were transduced with the DECIPHER 27K Pooled shRNA lentivirus library–Human Module 1 (Cellecta) at an MOI of 1.25 (based on 293T cell tittering) in the presence of polybrene and then spinoculated for 2 hours as described previously. The next day, half the cells were spun down and frozen as the *in vivo* baseline reference sample. The remainder of the cells were resuspended in 50% (v/v) mixtures of normal matrigel (BD Biosciences) and flow cytometry buffer in a total volume of 1.8 ml. These cells were injected evenly into the right and left mammary fat pads of seventeen NSG mice. When tumors reached approximately 10 mm in diameter, the mice were euthanized and the tumors dissected as previously described. These tumors were then processed into single cells, pooled, washed and frozen as the *in vivo* growth experimental sample.

The two pairs of samples, *in vitro* baseline reference sample and *in vitro* organoid growth experimental sample and *in vivo* baseline reference sample and *in vivo* growth experimental sample, were submitted to Cellecta, Inc. for genomic DNA extraction, bar code amplification, high-throughput sequencing and de-convolution. Twenty million barcode reads were performed for each sample.

### ‘Hit’ selection algorithm from the *in vivo* and *in vitro* RNAi dropout viability screens

Please see the schematic in Supplementary Figure 2A for an overview. We applied an algorithm to narrow our hits to a more manageable number for validation. 1) for each individual shRNA we determined a ‘dropout ratio’ that was shRNA barcode counts in the growth experimental sample divided by shRNA barcode counts in the baseline reference sample. In each screen, these were ranked from lowest to highest. 2) We examined the top 5% of the lowest dropout ratios in each experiment and identified genes targeted by ≥ 2 shRNA. 3) We cross-referenced the shRNA gene targets in the *in vivo* screen (208 genes) with those in the *in vitro* screen (150 genes) to identify genes that overlapped between the two experiments. These 46 overlapping ‘hit’ genes are shown in Supplementary Figure 2C.

### Single Cell Quantitative PCR Experiments

Sample preparation, single cell sorting, quantitative PCR and data analysis were as previously described^24^.

### Microarray Experiments

MDA-MB231 cells were infected with shCDK19-2, shCDK8-2 or control shRNA and grown in cell culture conditions for 96 hours. They were subsequently harvested by trypsinization, resuspended in flow cytometry buffer and sorted by flow cytometry to obtain cells that were both DAPI negative and RFP positive. RNA was extracted from these cells by Rneasy plus micro kit (Qiagen) according to manufacturer’s instructions and quantified on an Agilent 2100 Bioanalyzer. 50 ng of total RNA from each sample was used. *In vitro* transcription, fragmentation, labeling, hybridization to the microarray and scanning was performed by the Stanford Protein and Nucleic acid facility (PAN facility). Samples were hybridized on PrimeView Human Gene Expression Arrays (Affymetrix). Gene Level Differential Expression Analysis was performed with the Transcriptome Analysis Console (Affymetrix). Downregulated genes were defined as those for which log_2_ (sample/control) < −1.5 and upregulated genes log_2_ (sample/control) > 1.5.

### H3K27Ac Chromatin Immunoprecipitations

ChIP assays were performed as previously described ^47^. MDA-MB231 cells were infected with shCDK19-2, shCDK8-2 or control shRNA and grown in standard cell culture conditions for 96 hours before trypsinization for harvesting. Cells were sorted by FACS to identify DAPI negative and RFP positive cells that were the live cells transduced with shRNA. Approximately 250,000 to 500,000 MDA-MB231 cells were used per ChIP. 1μg of anti-H3K27ac (Active Motif #39133) were used per ChIP.

### Library construction

ChIP enriched DNA was quantified using a Qubit 3.0 and dsDNA HS assay. Up to 1 ng of DNA was used for library construction using transposition based NEXTERA XT (followed manufacturer’s protocol with ~14 PCR cycles for indexing). Indexed samples were pooled and submitted for sequencing on a NextSeq500 to obtain 75 bp single end reads with read depths of ~60 million reads.

### Sequence analysis

Raw sequence reads were uploaded to Galaxy (usegalaxy.org) and aligned to the human genome (hg19) using Bowtie2 (-very-fast-local). Only uniquely mapped reads were retained for further analysis (MAPQ > 1). To visualize data, alignment files were used to produce signal tracks with DeepTools (100 bp bins with 200 bp read extensions and RPKM normalization) and BigWig files were loaded into Broad’s Integrated Genome Browser. MACS2 was used to call peaks (-nomodel, p=0.001, -broad, 33utoff 0.1, duplicates = auto, extension 200) for each replicate. A consensus peak list containing only those peaks occurring in all replicates, was generated using Bedtools. We performed differential peak analysis across consensus peaks using DiffBind. The DiffBind output peak list was annotated by fetching the nearest non-overlapping feature of the human RefSeq table from UCSC. Data for aggregation plots of ChIP signal across various peaks sets were generated using DeepTools’ computeMatrix (scale-regions: 1000; 50 bp bins) and plotProfile. Data was then plotted with GraphPad Prism software.

### GSEA Analysis

Gene set enrichment analysis (GSEA) was performed using the javaGSEA desktop application (GSEA 3.0)^27^ with log_2_ fold change values for *CDK19* knockdown versus Control as the ranking metric and Hallmarks^28^, CDK19KD-EnhancerUp and CDK19KD-EnhancerDOWN as the gene sets that were tested for enrichment.

### Statistical Analysis

Results are shown as mean ± s.d. or mean mean ± s.e.m. Statistical calculations were performed with GraphPad Prism software (GraphPad Software Inc). Variance was analyzed using the F-test. To determine *P*-values, *t*-test was performed on homoscedastic populations, and *t*-test with Welch correction was applied on samples with different variances. For animal studies, sample size was not predetermined to ensure adequate power to detect a pre-specified effect size, no animals were excluded from analyses, experiments were not randomized and investigators were not blinded to group allocation during experiments.

### Data Availability

Raw and final processed data related to this paper have been deposited in GEO (https://www.ncbi.nlm.nih.gov/geo/). The CHIP-Seq data has been deposited with an accession number of GSE103887 and the microarray data has been deposited with an accession number of GSE103851.

## Acknowledgements

We would like to thank Dr. Steve Chan for advice regarding CDK19 validation, Dr. Roel Nusse for the gift of L1-Wnt3a feeder cells, Pauline Chu for assistance with sample preparation, Patty Lovelace and Jennifer Ho for assistance with flow cytometry, Natalia Kosovilka and the Stanford PAN facility for assistance with microarray experiments, Gunsagar Gulati for single cell qPCR interpretation and Gary Mantalas and Dr. Norma Neff for technical consultations regarding gene expression.

This work was supported by NIH/NCI 5R01 CA100225, Department of Defense grant W81XWH-11-1-0287, Department of Defense/Breast Cancer Research Program (BCRP) Innovator Award W81XWH-13-1-0281, The Breast Cancer Research Foundation and Ludwig Cancer Research. The flow cytometry work was supported by NIH S10 Share Instrument Grant (1S10RR02933801). RWH was supported by the California Institute of Regenerative Medicine Postdoctoral Fellowship Training Grant and the Stanford Cancer Institute 2015 Fellowship Award Grant. The authors declare no competing financial interests.

## Author Contributions

RWH, AHK, FAS, MAZ and MFC designed the research and analyzed the data. All experiments were performed under guidance from MFC. Specifically, RWH performed CDK19 *in vitro* and *in vivo* validation studies, AHK and FAS performed tumor initiating cell identification and limiting dilution experiments, RWH and MAZ performed CHIP-Seq experiments, RWH performed microarray studies and GSEA, RWH performed *in vivo* pharmacologic inhibition experiments, SSS, SS and TK performed single cell quantitative PCR experiments, JA and LSH performed fluorescence microscopy, RWH and DP performed molecular cloning and RWH and AHK performed quantitative PCR studies. AHK and SSS advised on *in vivo* and cellular *in vitro* studies. AHK, SSS, FAS and DQ prepared patient derived xenografts. GS and FMD obtained patient samples from which PDXs were derived. SM and AJ provided compounds and input into the small molecule inhibition experiments. AMN assisted with microarray data processing and advised on the GSEA, Microarray and CHIP-Seq analysis. RWH and MFC wrote the manuscript with editing provided by AHK, FAS, MAZ, SSS and AMN.

## Supplementary Tables

**Supplementary Table 2. List of all CDK19KD-EnhancerUp and CDK19KD-EnhancerDOWN genes. (A and B)** There are (A) 1593 CDK19KD-EnhancerUp genes and (B) 341 CDK19KD-EnhancerDOWN genes. Highlighted in yellow are the (A) 478 CDK19KD-EnhancerUp ‘core’ and (B) 119 CDK19KD-EnhancerDOWN ‘core’ genes. Rank in Gene List, Rank Metric Scores and Running Enrichment Scores as determined by GSEA are also shown for each gene.

## Supplementary Figure 1

**Supplementary Figure 1.**
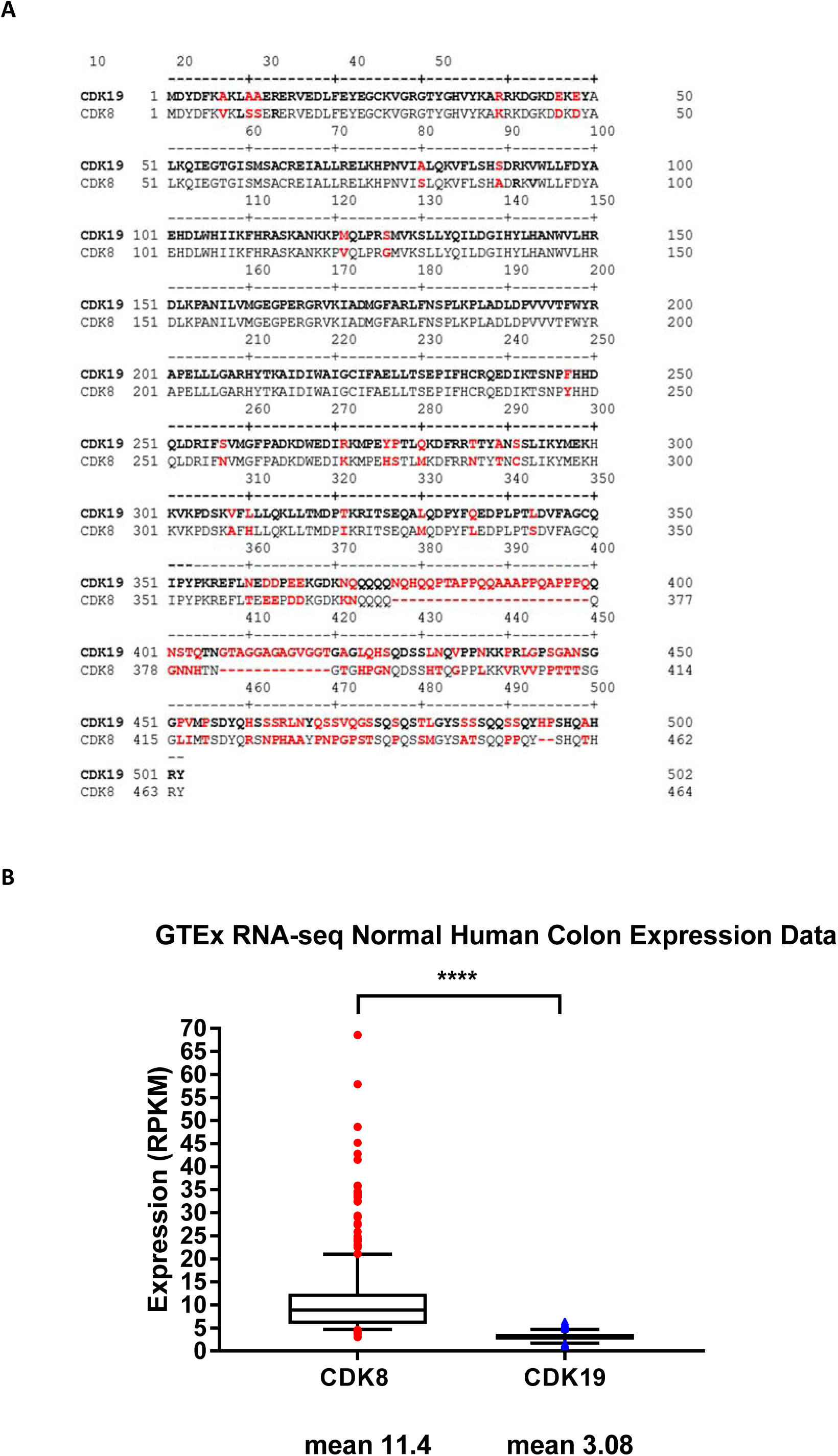
(A) **CDK19 and CDK8 share significant sequence homology** Amino acid sequence alignments show 84% sequence homology between CDK19 and CDK8. The amino acid differences between CDK19 (bold font, upper sequence) and CDK8 (normal font, lower sequence) are highlighted in red. The vast majority of differences between CDK19 and CDK8 are in the C-terminal domain. Amino acid positions are shown above the sequence. Alignment performed using Clustal W method with MegAlign (DNAStar).
(B) **Expression of *CDK8* and *CDK19* in normal human colon from the GTEx database.** Shown are box and whisker plots of *CDK8* and *CDK19* expression as determined by RNA-Seq in the GTEx database of human normal tissue (mean shown, n = 345) (Line in Box shows the median, while box lower and upper edges show 25^th^ and 75^th^ percentiles respectively. Error bars represent 10^th^ - 90^th^ percentiles).

## Supplementary Figure 2

**Supplementary Figure 2.**
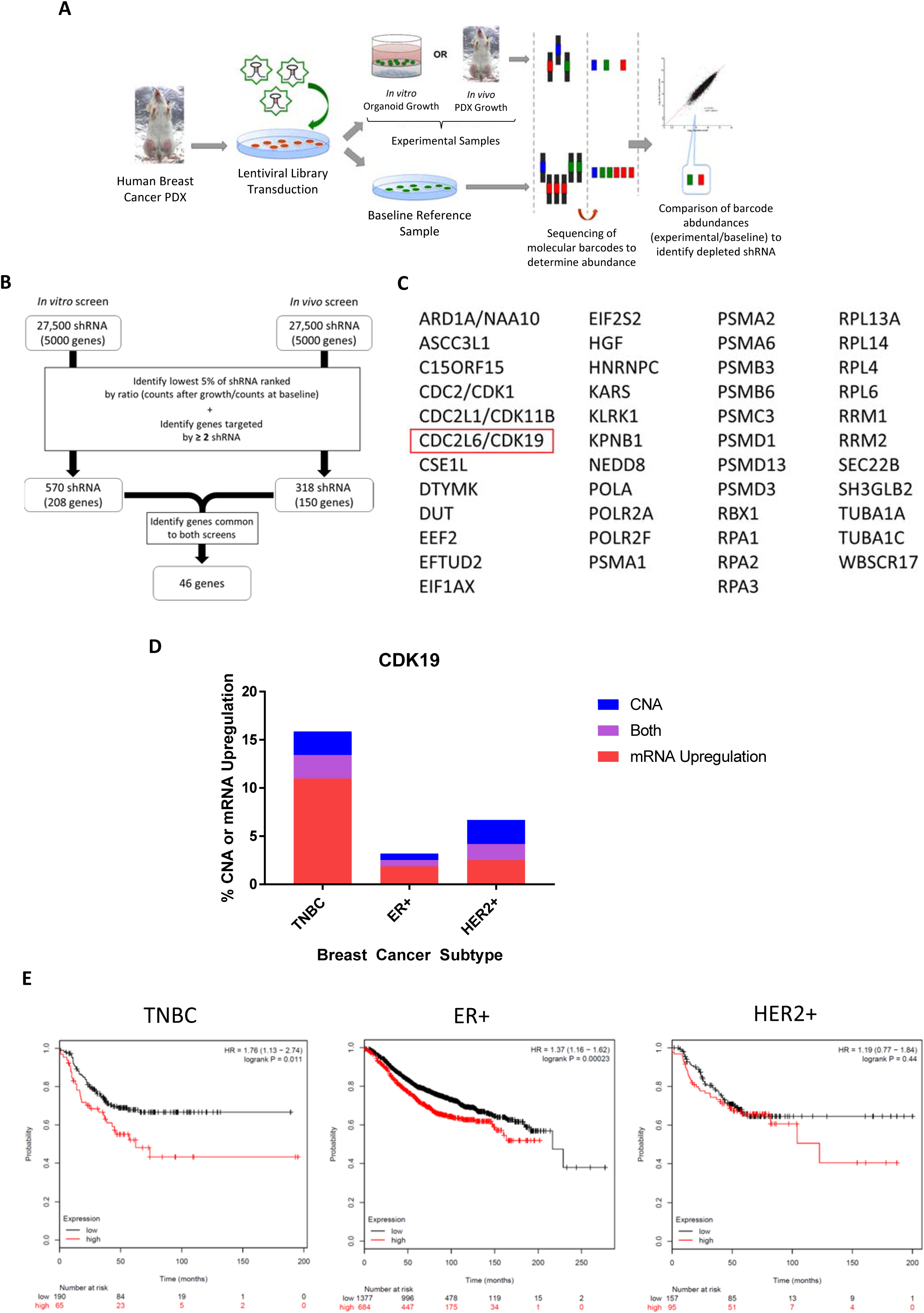
Selection of CDK19 as candidate gene from in vitro and in vivo RNAi screens. (A) **Schematic for the RNAi dropout viability screens**. Two separate screens were performed in a TNBC PDX (PDX-T1). Cells in one experiment were grown in vitro as organoid colonies and in the other in vivo as PDXs in NSG mice. The abundance of individual shRNA in each experimental sample and the baseline reference samples were determined by high throughput sequencing of the shRNA barcodes. The barcode counts in the experimental samples were compared to counts in the baseline reference sample to identify depleted or enriched barcodes.
(B) Schematic of the criteria used to narrow the initial list of hits from the in vitro and the in vivo screens down to 46 candidate genes.
(C) List of 46 candidate genes determined from the in vitro and the in vivo screens after filtering with the criteria shown in (B). CDK19 is highlighted (red box).
(D) TCGA breast cancer samples from patients with the TNBC subtype are enriched in CDK19 copy number amplifications and/or CDK19 mRNA upregulation compared to other subtypes. Percentage of samples with CDK19 copy number amplifications or CDK19 mRNA upregulation in triple-negative (TNBC), HER2 positive (HER2+) and estrogen receptor positive (ER+) breast cancers. Data obtained from cBioPortal and based on TCGA data.
(E) Relapse free survival for different breast cancer subtypes stratified by CDK19 expression. Figures generated using an online tool (Kaplan-Meier Plotter) based on microarray data.

## Supplementary Figure 3

**Supplementary Figure 3.**
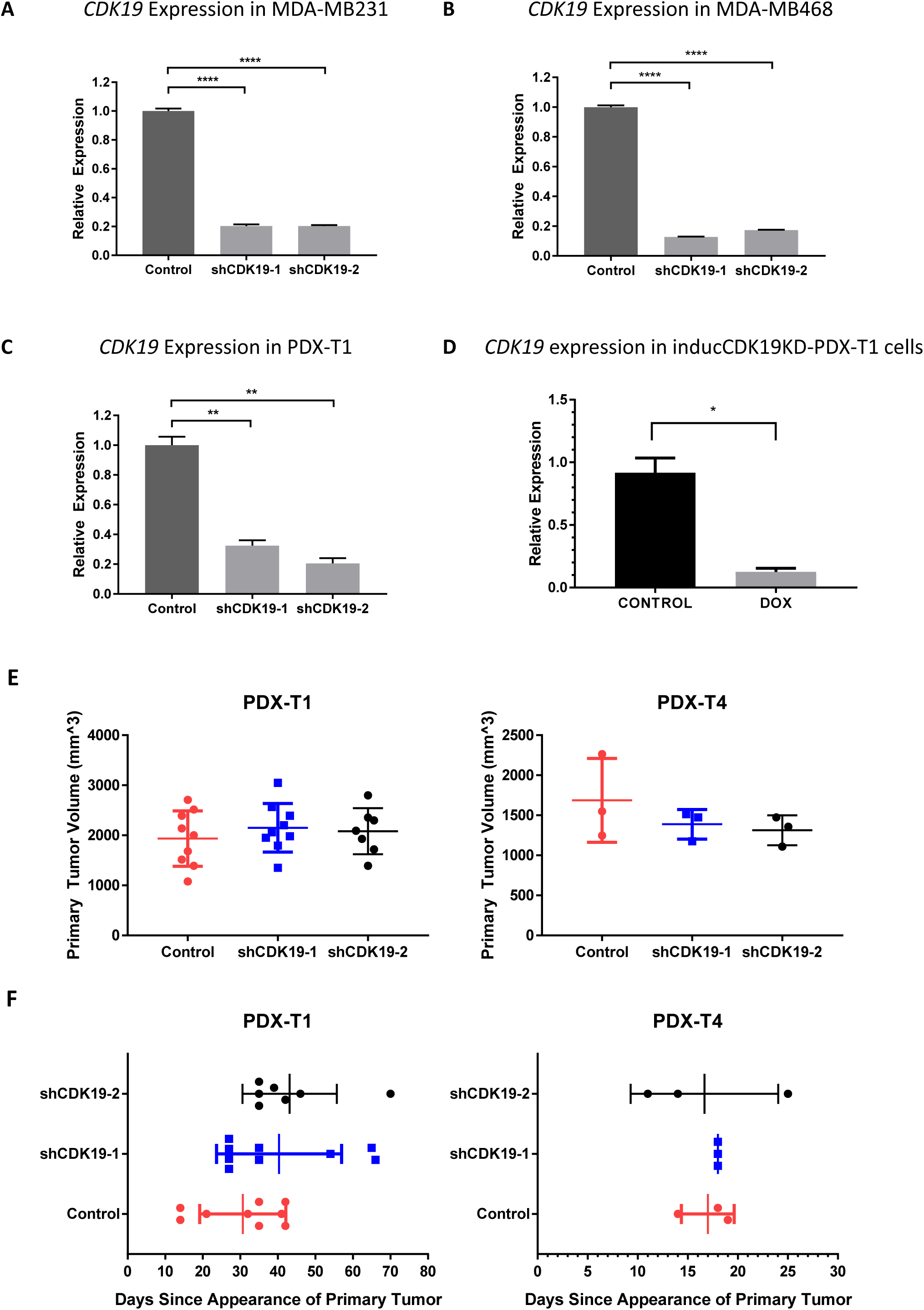
The expression of *CDK19* in cell lines and PDXs in response to *CDK19* knockdown. (A and B) Expression of *CDK19* in (A) MDA-MB231 or (B) MDA-MB468 determined by RT-qPCR for cells transduced with control shRNA, shCDK19-1 and shCDK19-2 (n = 3, experiments performed twice).
(C) Expression of *CDK19* in PDX-T1 as determined by RT-qPCR for cells transduced with control shRNA, shCDK19-1 and shCDK19-2 (n = 2, experiments performed twice).
(D) Expression of *CDK19* in inducCDK19-PDX-T1 with doxycycline treatment (DOX) and without doxycycline (CONTROL) as determined by RT-qPCR (n = 2, experiments performed twice). For all, the relative expression of *CDK19* in *CDK19* knockdown cells is normalized to the mean expression of *CDK19* in cells transduced with control shRNA. Gene expression in each condition is normalized to beta-actin as a housekeeping gene (mean + s.d., \**P* < 0.05, \*\**P* < 0.01; \*\*\*\**P* < 0.0001)
(E and F) (E) Primary tumor volumes and (F) days since appearance of primary tumor at the time that mice were examined for metastasis are shown for the tumors treated with control or CDK19 shRNA in PDX-T1 (left) and PDX-T4 (right). Data are represented as mean ± s.d. No statistically significant differences are noted between any of the treatment conditions in any of the experiments (one-way ANOVA).

## Supplementary Figure 4

**Supplementary Figure 4.**
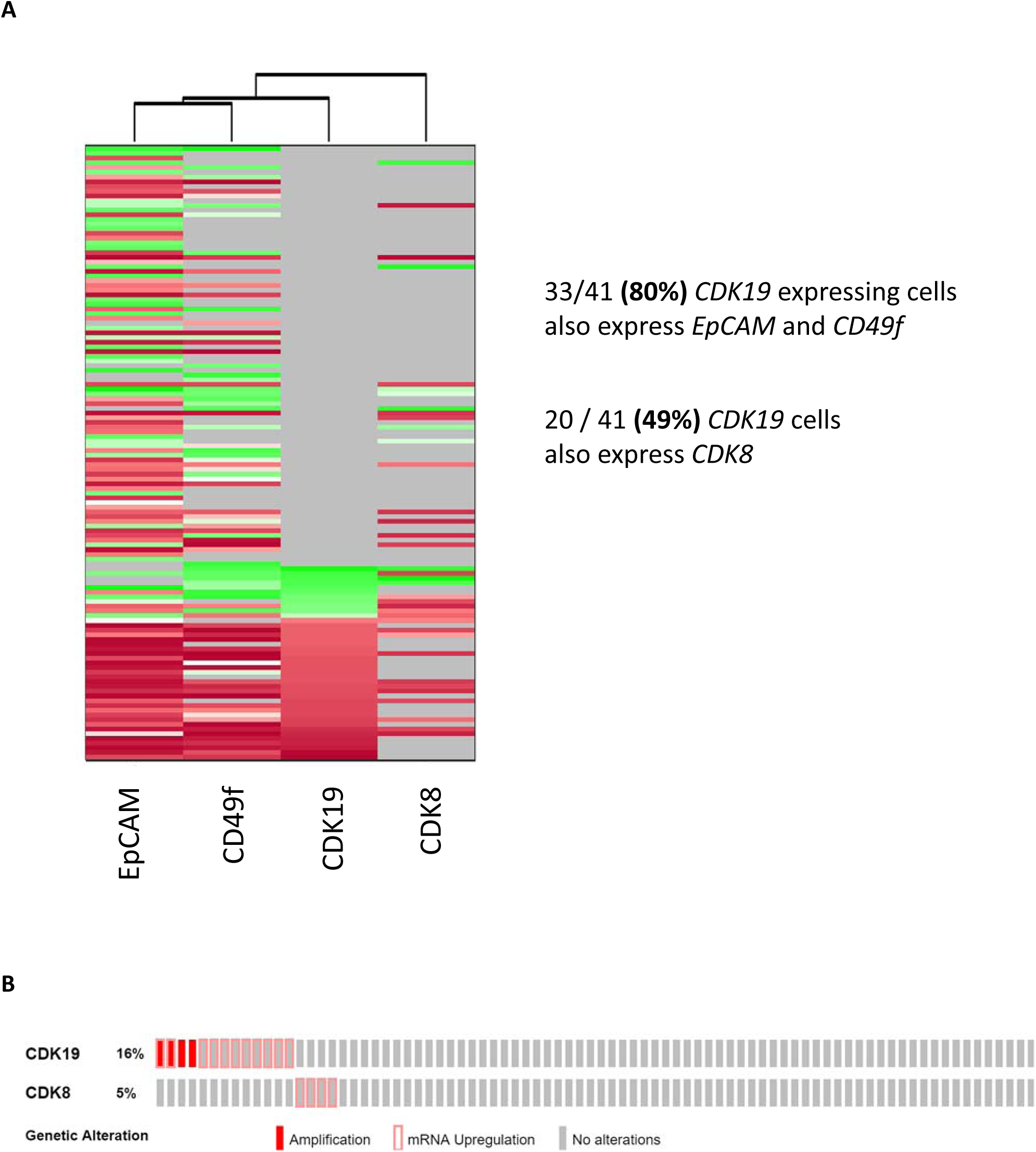
(A) ***EpCAM and CD49f* are co-expressed in *CDK19* expressing cells.** Single cell quantitative PCR gene expression heatmap of primary TNBC cells from which PDX-T1 was derived. Single cells (rows) and genes (columns) were measured simultaneously from each cell. Cells were sorted from top to bottom by increasing expression of *CDK19*. Genes were clustered such that phenotypically similar cells were grouped next to each other (Red – high expression, Green – low expression, Gray – no expression).
(B) **CDK19 mRNA upregulation and/or copy number amplifications is mutually exclusive of CDK8 mRNA upregulation and/or copy number amplifications in patient tumors.** CDK19 and CDK8 mRNA upregulation (empty red box) or copy number amplifications (filled red box) in triple-negative breast cancer patient tumors from the TCGA. Each column represents one patient tumor. There were 82 patient tumors with TNBC from the data set (not all patient tumors are shown. Patient tumors not shown had no alterations).

## Supplementary Figure 5

**Supplementary Figure 5.**
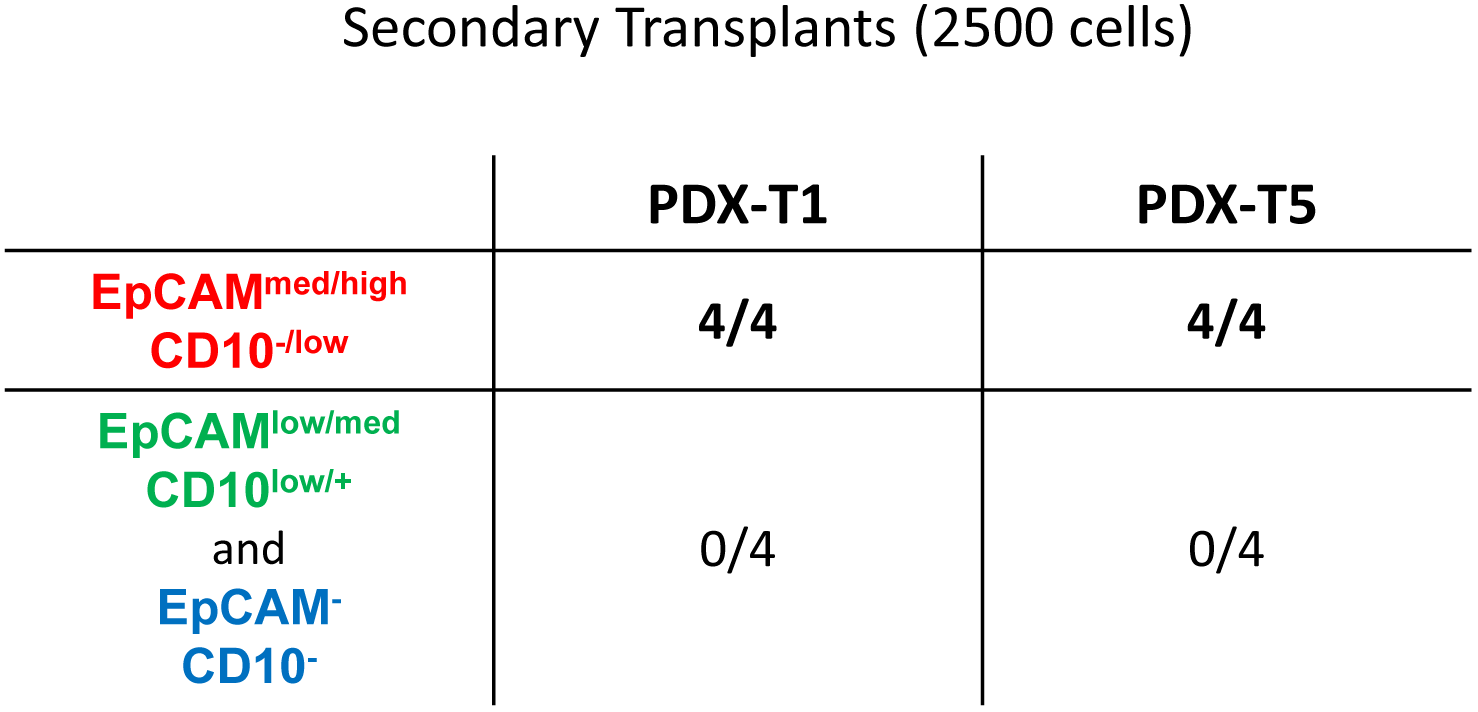
Secondary Transplants of PDXs to evaluate Tumor Initiating Capacity. For the secondary transplants, dissociated PDX tumor cells from the primary transplant were sorted into sub-populations based on the protein expression of EpCAM and CD10. 2500 cells were then injected into the mammary fat pads of NSG mice. The number of tumors formed and the number of injections performed are indicated for each population. Populations and injections where tumors formed are bolded.

## Supplementary Figure 6

**Supplementary Figure 6.**
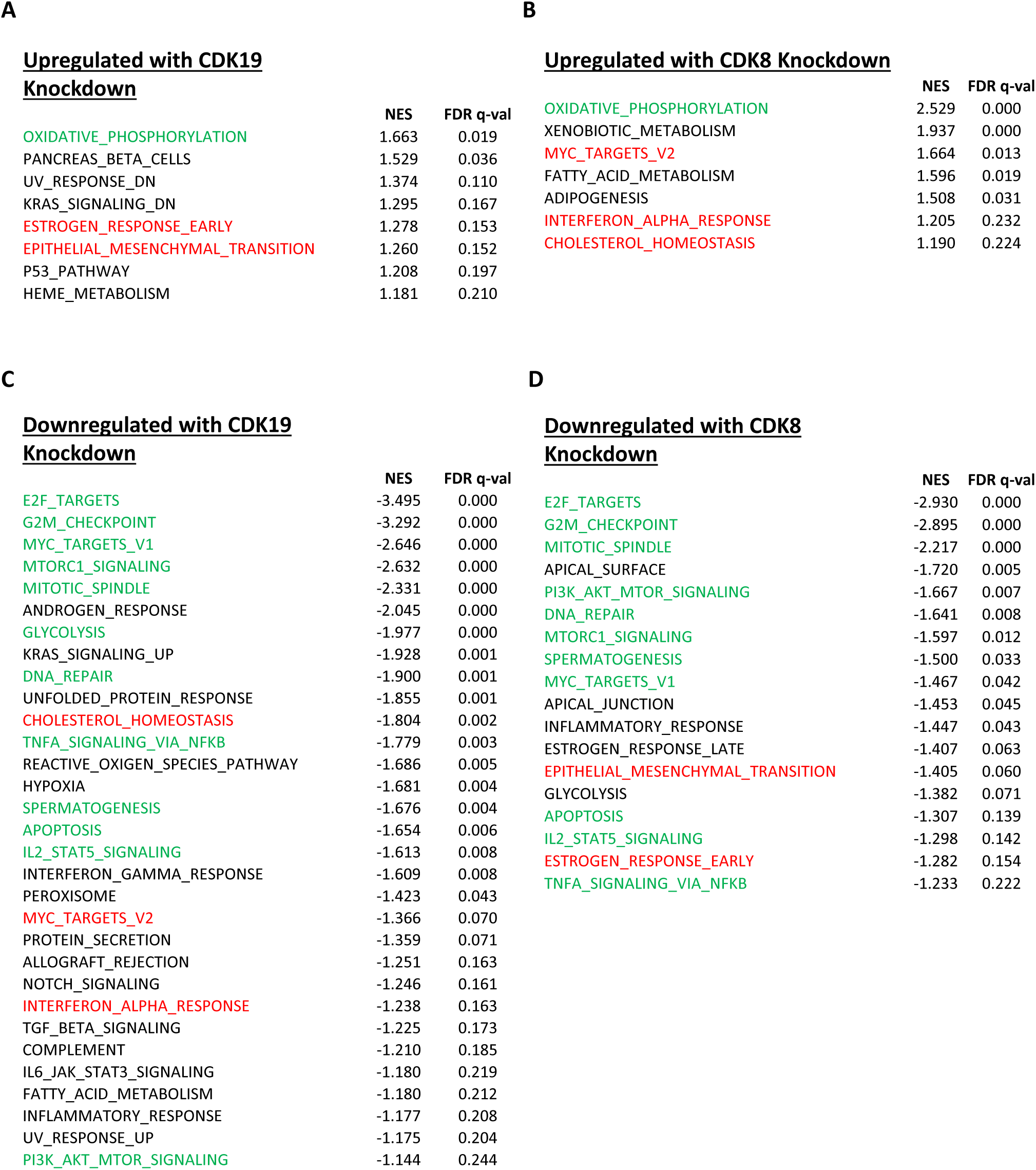
Hallmark gene sets enriched by *CDK19* knockdown and *CDK8* knockdown. **(A, B, C, D)** Hallmark gene sets found enriched by GSEA of the genes upregulated or downregulated by either (A and C) *CDK19* knockdown or (B and D) *CDK8* knockdown. The Hallmark gene sets uniquely enriched in the knockdown of *CDK19* or *CDK8* are shown in black, enriched in both the knockdown of *CDK19* and *CDK8* are shown in green and enriched by genes expressed in opposite directions between the knockdown of *CDK19* and *CDK8* are shown in red. Normalized enrichment scores and FDR q-values are determined by the GSEA software. An FDR q-value cutoff of < 0.25 was used to select significant gene sets.

## Supplementary Figure 7

**Supplementary Figure 7.**
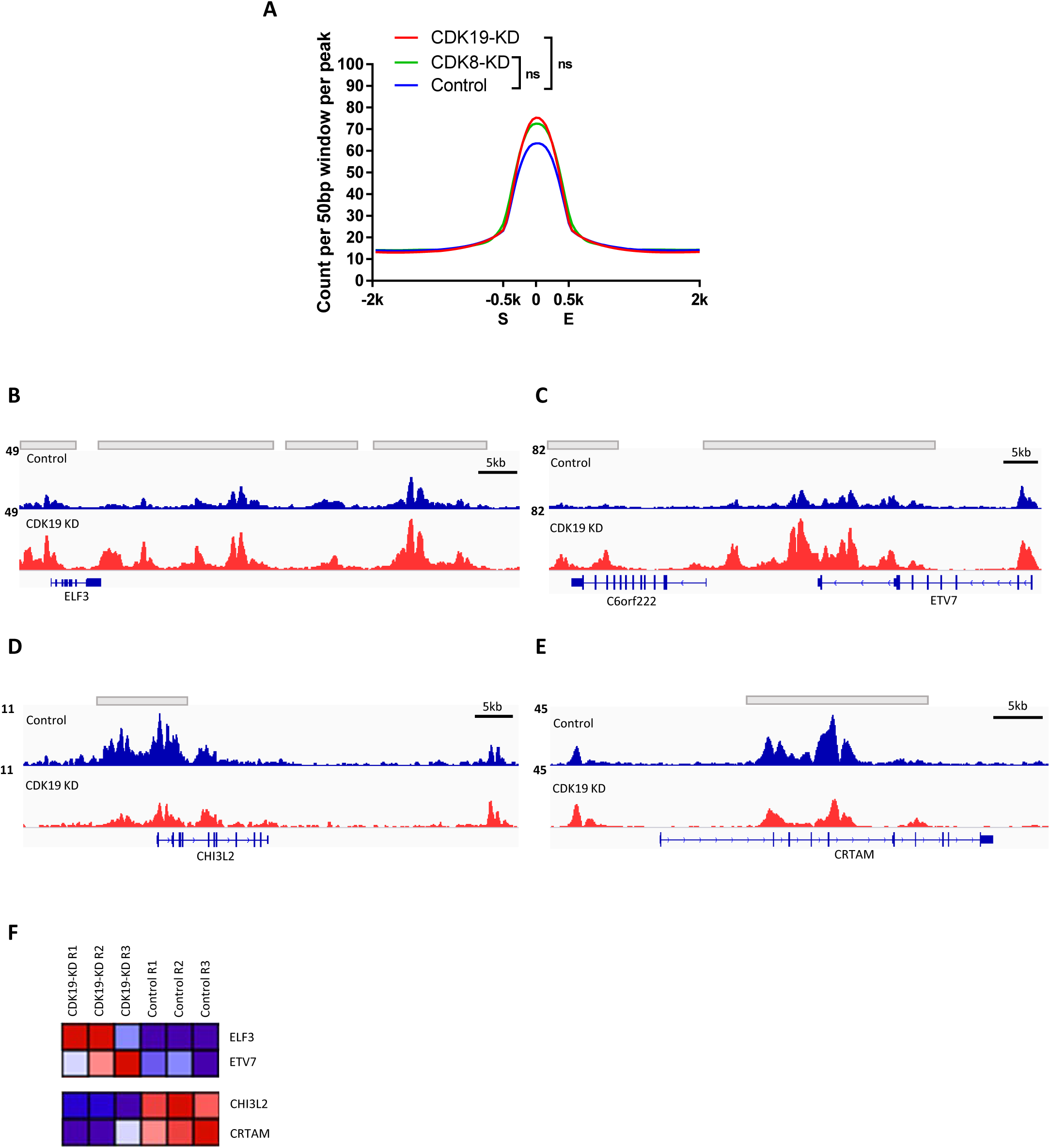
Genome-wide H3K27Ac CHIP-Seq signal assessment and representative genes where changes in H3K27Ac signals correspond strongly to gene expression. (A) Genome-wide H3K27Ac CHIP-Seq signals across *all* identified H3K27Ac regions. Aggregate plots of normalized H3K27Ac CHIP-Seq signals across all H3K27Ac regions in the *CDK19* knockdown (blue line), *CDK8* knockdown (green line) and control (red line) samples (ns is *P* > 0.05, all samples n = 3, experiments performed three times). H3K27Ac CHIP-Seq signals across all identified H3K27Ac regions are normalized to 1-Kb and centered on the middle of those regions. Signals of the flanking 2-Kb regions are also shown. To compare relative signal changes, the total signal of each biological replicate was determined by summing the signals of each 50-base window 1-Kb around the center of each region (S and E denote start and end, respectively). *P*-values between total CHIP-Seq signals of each sample were determined by unpaired t-test.
(B-E) Representative gene tracks depicting H3K27Ac signals at the loci of **(B and C)** CDK19KD-EnhancerUP and **(D and E)** CDK19KD-EnhancerDOWN genes. In comparing the control and *CDK19* knockdown gene tracks, the gene tracks at the (B) *ELF3* and (C) *ETV7* loci show relative enrichment of H3K27Ac signals in the *CDK19* knockdown samples, whereas the gene tracks at the (D) *CHI3L2* and (E) *CRTAM* loci show relative enrichment of H3K27Ac signals in the Control samples. Upper tracks denote Control samples (blue), while lower tracks denote *CDK19* knockdown samples (red). Grey bars denote regions identified by DiffBind to be different between control and *CDK19* knockdown samples (FDR < 0.05).
(F) Heat map of the normalized gene expression of *ELF3*, *ETV7*, *CHI3L2* and *CRTAM* across each of the three biological replicates in control and *CDK19* knockdown samples (Red – high expression, Blue – low expression).

## Supplementary Figure 8

**Supplementary Figure 8.**
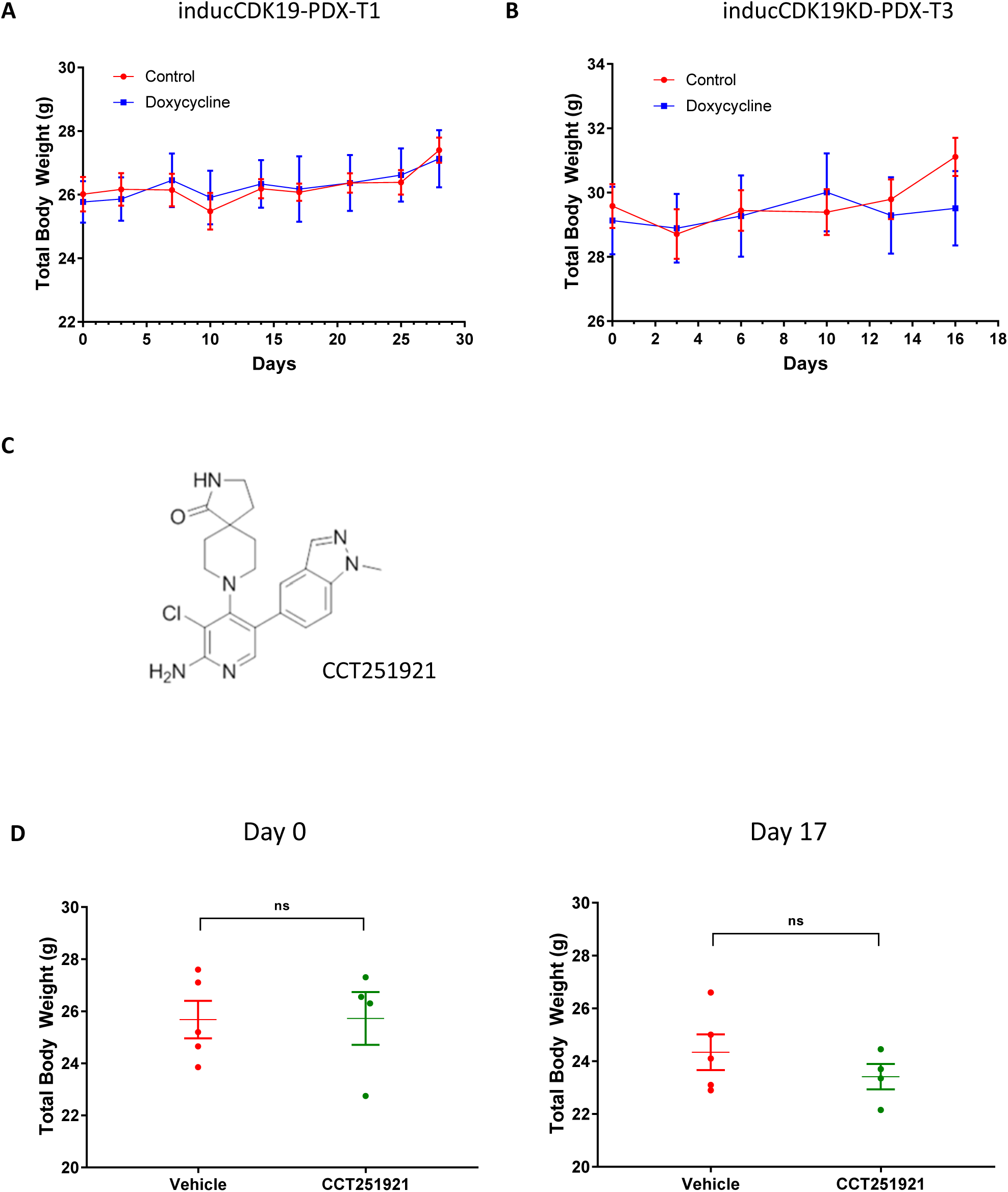
Genetic knockdown of *CDK19* or chemical inhibition of CDK19 kinase activity did not affect weights relative to the weights of control mice. (A and B) Total body weights of mice were not significantly different between the mice fed doxycycline rodent feed (doxycycline group, blue) compared to the mice fed standard rodent feed (control group, red) in the (A) inducCDK19KD-PDX-T1 (mean ± s.e.m, 10 mice for each condition tested) and (B) inducCDK19KD-PDX-T3 (mean ± s.e.m, 10 mice for each condition tested) tumor experiments.
(C) Chemical structure of CCT251921 that was used in the chemical inhibition experiments.
(D) Total body weights of mice were not significantly different between the mice receiving oral gavage with CCT251921 (green) compared to Vehicle (red) at the beginning of the experiment on day 0 (left) and at the end of the experiment on day 17 (right) (mean ± s.e.m, 4 −5 mice total in each treatment condition).

## Supplementary Table 1

**Supplementary Table 1.** Pathological features and patient information for the patient derived xenograft tumors used in this manuscript.

| **Tumor** | **Origin** | **ER/PR/HER2**<br>**status** | **Diagnosis** | **Treatment**<br>**Status** |
| --- | --- | --- | --- | --- |
| PDX-T1 | Breast<br>Primary | -/-/- | IDC | Radiation |
| PDX-T2 | Breast<br>Primary | -/-/- | IDC | Not treated |
| PDX-T3 | Breast<br>Primary | -/-/- | IDC | Not treated |
| PDX-T4 | Brain<br>metastases | -/-/- | Inflammatory<br>Breast CA | Neo-adjuvant<br>chemotherapy |
| PDX-T5 | Breast<br>Primary | -/-/+ | Inflammatory<br>Breast CA | Neo-adjuvant<br>chemotherapy |
| PDX-T6 | Breast<br>Primary | -/-/- | IDC | Neo-adjuvant<br>chemotherapy |
| PDX-T7 | Breast<br>Primary | +/+/+ | IDC | Unknown |
| PDX-T8 | Breast<br>Primary | +/+/- | IDC | Neo-adjuvant<br>chemotherapy |

